# Intragenic redistribution of RNA polymerase II licenses epigenetic silencing

**DOI:** 10.64898/2026.11.19.766859

**Authors:** Xin Song, Yujuan Liu, Liangbing Yuan, Yuantao Hu, Jingjing Li, Yixiong Zhong, Hongyu Huo, Yisui Huang, Zhihao Zhang, Zhenwei Liang, Tao Zhu, Chen Chen, Chenlong Li

## Abstract

Co-transcriptional chromatin regulation is widely recognized as a mechanism that reinforces active gene expression, yet transcriptionally active genes must also be able to transition into stable epigenetic silencing during developmental transitions and environmental responses. Whether and how ongoing transcription is coupled to the initiation of epigenetic silencing remains unclear. Here we show that elongating RNA polymerase II phosphorylated at Ser2 (RNAPII-Ser2P) functions as an instructive signal for epigenetic silencing initiation. At actively transcribed genes, RNAPII-Ser2P positions and spatially constrains the FLD–LD H3K4me1 demethylase complex near transcription termination regions, separating the complex from its gene-body H3K4me1 substrate. We found that both chemical perturbation of transcription and physiological silencing cues induce a redistribution of RNAPII-Ser2P into the gene body, accompanied by FLD relocalization and subsequent gene-body H3K4me1 erasure. FLD disengages from chromatin upon gene silencing, indicating that active transcription is required for its continued chromatin association. We further show that LD tethers FLD to RNAPII through an N-terminal HOX domain that binds the C-terminal region of FLD and an adjacent TFIIS domain that contacts the RNAPII CTD in a Ser2P-sensitive manner. This FLD–LD–RNAPII bridge is essential to promote H3K4me1 erasure and gene silencing. Together, these findings reveal a mechanism by which active transcription licenses the initiation of epigenetic silencing.

## Main Text

A hallmark of developmental and cell fate transitions is the stable activation of genes that were previously silent. Gene activation is typically accompanied by characteristic chromatin changes^1-3^, including increased accessibility^4-6^, histone acetylation^7-10^, promoter-proximal H3K4 trimethylation^11-17^ and gene-body H3K36 methylation^18-24^, all of which support transcriptional competence. RNA polymerase II (RNAPII) is central to this coupling: as it progresses through transcription initiation and elongation, it coordinates the recruitment of diverse chromatin regulators that further strengthen the transcriptionally active state, aided partly by dynamic phosphorylation of its carboxyl-terminal domain (CTD)^25-29^. Together, these mechanisms establish that transcription is not merely an output of gene activation but also a reinforcing force that helps stabilize and maintain active chromatin states^9,27,30^.

Developmental and cell fate transitions, however, also require genes to become silenced from a state of ongoing transcription^31-37^. This creates a fundamental paradox: the same transcription machinery that promotes active chromatin states must somehow be co-opted to initiate the opposite transition. Many existing models resolve this paradox indirectly by invoking the RNA products as instructive intermediates; for example, siRNAs nucleate heterochromatin^38,39^, piRNAs enforce transposon silencing^40-43^, and lncRNAs such as *Xist* concentrate Polycomb activity at target loci^44,45^. In these frameworks, the silencing-instructive function of transcription is conveyed principally by the RNA molecules, leaving RNAPII itself in a supporting role. However, RNA-centered models leave unresolved how a newly synthesized RNA determines that a locus should undergo epigenetic repression rather than remain active and, more fundamentally, how repressive chromatin regulators are selectively targeted to particular transcribed loci despite pervasive transcription across the genome. Whether RNAPII itself, beyond the RNAs it generates, can provide instructive information for epigenetic silencing remains unclear.

If RNAPII contributes to silencing decisions directly, an important question is what feature of the transcription machinery encodes such information. Elongating RNAPII does not progress through gene bodies in a uniform manner^46^. Its movement across gene bodies is shaped by pausing^47-50^, chromatin architecture^51^ and regulatory inputs^52-54^, generating substantial variation in the local occupancy of RNAPII across individual loci. Together, these local differences create distinct co-transcriptional environments within a single gene^55-57^. Whether these intragenic RNAPII occupancy states can specify where and when repressive chromatin transitions are initiated remains unknown.

A particularly informative system for addressing this question is the removal of H3K4 monomethylation (H3K4me1), which mediates the transition from active transcription to epigenetic silencing. H3K4me1 is a conserved histone modification associated with transcriptionally active chromatin across eukaryotes^58-60^. Although its genomic distribution varies among species, H3K4me1 in plants is predominantly enriched across gene bodies, in contrast to H3K4 trimethylation (H3K4me3) and H3K4me2, which accumulate near transcription start sites (TSSs)^11,61-63^. Genetic and molecular studies in plants have established that removal of gene-body H3K4me1 is critical for the transition from transcriptionally active to silent chromatin states^64-67^. However, whether and how gene-body H3K4me1 erasure is coupled to ongoing transcription to initiate gene silencing remains unclear.

Here we show that elongating RNAPII positions an H3K4me1 demethylase complex on actively transcribed chromatin, thereby coupling active transcription to a silencing-permissive chromatin transition. At actively transcribed genes, Ser2-phosphorylated RNAPII (RNAPII-Ser2P) spatially restricts this demethylase complex near transcription termination regions, limiting its access to H3K4me1-rich gene bodies. Using a tractable silencing model, we show that chemical perturbation of transcription or physiological silencing cues induce redistribution of RNAPII-Ser2P and the H3K4me1 demethylase complex into the gene body, relieving this spatial constraint and enabling subsequent gene-body H3K4me1 erasure during epigenetic silencing initiation. Our findings identify elongating RNAPII as a chromatin-state decision point, revealing how the same transcription machinery that sustains an active state can also license the transition into epigenetic silencing.

## Results

### Elongating RNAPII-Ser2P positions FLD on chromatin

To investigate how active transcription may be coupled to the initiation of epigenetic silencing, we examined the chromatin engagement of the H3K4me1 demethylase FLOWERING LOCUS D (FLD), a key effector of gene silencing in plants^68-72^. We performed ChIP-seq for FLD in *Arabidopsis* seedlings expressing a *pFLD:FLAG-GFP-FLD* transgene (FG-FLD), and profiled Ser2-phosphorylated RNA polymerase II (RNAPII-Ser2P) in matched seedlings. RNAPII-Ser2P marks elongating RNAPII and preferentially accumulates toward transcription termination sites (TTSs) at actively transcribed genes^69,73-75^. Consistent with previous work^69^, FLD occupancy significantly overlapped with RNAPII-Ser2P-enriched loci genome-wide, with 5,368 of 8,816 RNAPII-Ser2P target genes also occupied by FLD (Fig. 1a-c).

**Fig. 1.**
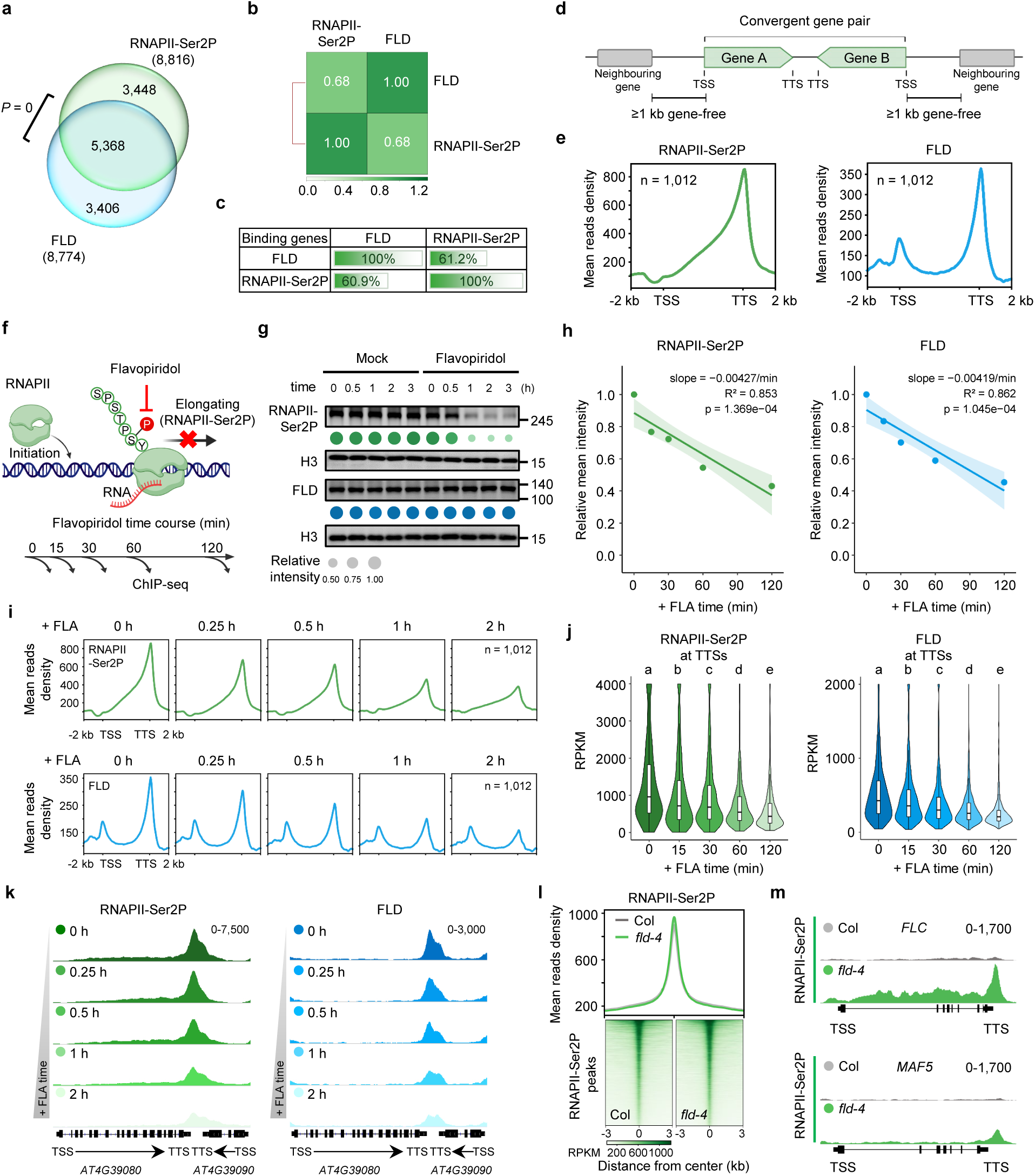
Elongating RNAPII-Ser2P positions FLD on chromatin. **a**, Venn diagram illustrating the statistically significant overlap between genes occupied by RNAPII-Ser2P (8,816 target genes) and FLD (8,774 target genes), of which 5,368 genes are co-occupied by both. The *P* value was calculated using the hypergeometric test. **b**, Matrix depicting correlation coefficients between ChIP-seq datasets. **c**, Percentages of FLD- and RNAPII-Ser2P-bound genes (by row) overlapping with other bound genes (by column). **d**, Schematic of the convergently transcribed genes used in this study. A convergent gene pair comprises two genes oriented towards each other, such that their transcription termination sites (TTSs) face one another, placing a termination region rather than a promoter immediately 3′ of each TTS. Gene pairs were retained only when ≥1 kb outside each transcription start site (TSS) was free of annotated genes on either strand, so that TSS-proximal signal was separated from the nearest neighbouring locus. No constraint was placed on the distance between the two facing TTSs. Both genes of a retained pair were counted separately in n. These criteria were met by 1,012 of the 8,816 RNAPII-Ser2P target genes. **e**, Metagene plots displaying RNAPII-Ser2P and FLD ChIP-seq signals at RNAPII-Ser2P target genes that meet the criteria in **d** (*n* = 1,012). TSS, transcription start site; TTS, transcription termination site. **f**, Schematic of the experimental design for RNAPII elongation inhibition by flavopiridol treatment for the indicated times. **g**, Immunoblot analysis showing RNAPII-Ser2P and FLD protein levels in seedlings after treatment with flavopiridol for the indicated times. Dot sizes below indicate amounts normalized to histone H3. **h**, Kinetic changes in the relative mean intensity of RNAPII-Ser2P and FLD chromatin occupancy after flavopiridol treatment. Analyses were performed on RNAPII-Ser2P target genes meeting the criteria in **d** (*n* = 1,012; the same gene set was used in **i**–**k**. Shaded areas represent 95% confidence intervals. **i**, Metagene plots displaying the ChIP-seq signals of RNAPII-Ser2P and FLD at indicated time points after flavopiridol treatment. **j**, Violin plots showing RNAPII-Ser2P and FLD ChIP-seq read counts within ±500 bp of the TTS after flavopiridol treatment. Different letters indicate statistically significant differences. **k**, Integrative Genomics Viewer (IGV) tracks of RNAPII-Ser2P and FLD ChIP-seq signals at representative RNAPII-Ser2P target genes after flavopiridol treatment. Black diagrams underneath indicate gene structures. **l**, Genome-wide comparison of RNAPII-Ser2P binding profiles between wild-type Col and *fld-4* mutant backgrounds. Top: metagene plots displaying RNAPII-Ser2P ChIP-seq signals at binding peaks across the genome. Bottom: heatmap of RNAPII-Ser2P ChIP-seq signals at the same binding sites for each genotype. **m**, IGV tracks of RNAPII-Ser2P ChIP-seq reads at the *FLC* and *MAF5* loci in Col and *fld-4*.

Because the compact *Arabidopsis* genome places many genes in close proximity^69,76^, ChIP-seq signals from neighbouring loci can confound the spatial distribution of RNAPII-Ser2P and FLD around individual genes. We therefore examined convergently oriented RNAPII-Ser2P target genes, whose immediately downstream neighbour lies on the opposite strand and is transcribed towards them (n = 1,012; Fig. 1d and Supplementary Table 1). At these genes, FLD was preferentially enriched near the TTSs, where RNAPII-Ser2P also accumulated (Fig. 1e), consistent with the reported TTS-proximal colocalization of FLD with RNAPII-Ser2P on actively transcribed chromatin^69^.

To test whether RNAPII-Ser2P is required to sustain FLD chromatin association, we used fast-acting transcription elongation inhibitor flavopiridol to acutely inhibit RNAPII Ser2 phosphorylation (Fig. 1f,g). Flavopiridol treatment caused a rapid reduction in cellular RNAPII-Ser2P levels (Fig. 1g)^75,77,78^, and in line with this, RNAPII-Ser2P ChIP-seq signals progressively declined genome-wide, dropping by ∼50% within 1 h and remaining reduced at 2 h (Fig. 1h-k and Extended Data Fig. 1a). As expected, the decrease in RNAPII-Ser2P enrichment was found primarily at the TTSs (Fig. 1i-k and Extended Data Fig. 1c,d). Strikingly, FLD chromatin occupancy rapidly decreased at the same TTS-proximal regions with closely matched kinetics across the genome (Fig. 1h-k and Extended Data Fig. 1b-d), despite unchanged FLD protein abundance (Fig. 1g), indicating that RNAPII-Ser2P is required to sustain FLD association with TTS regions at actively transcribed genes.

We asked whether this relationship was reciprocal. Loss of FLD did not reduce genome-wide RNAPII-Ser2P occupancy (Fig. 1l). Notable exceptions included the flowering regulators *FLOWERING LOCUS C* (*FLC*) and its related MADS-box gene *MADS AFFECTING FLOWERING 5* (*MAF5*), where RNAPII-Ser2P enrichment levels were elevated in *fld*, consistent with their transcriptional de-repression in this background (Fig. 1m). Thus, RNAPII-Ser2P promotes FLD chromatin association, whereas FLD is generally not required for RNAPII-Ser2P occupancy.

Previous work showed that FLD acts genetically downstream of FLOWERING CONTROL LOCUS A (FCA) and the SPEN-family protein FPA, two RNA-binding proteins that promote proximal transcription termination on both strands of *FLC*.^70,72,79-82^ Interestingly, FLD chromatin occupancy was not reduced genome-wide in *fca* or *fpa* mutants (Extended Data Figs. 2 and 3a-f), despite disrupted co-transcriptional RNA processing in these mutants^71,72,80,83^. At *FLC*, whose transcription is elevated in *fca* and *fpa*, FLD occupancy instead increased at the TTS region (Extended Data Fig. 3g-j)^72^, consistent with transcription-coupled FLD recruitment. Reciprocally, in the *FPA-GFP fpa* line, loss of FLD did not reduce FPA occupancy genome-wide at FPA target genes and around FPA-bound TTS regions (Extended Data Fig. 3k-n). Notably, paralleling the rise in *FLC* transcription in the *fld* mutant, FPA occupancy at the de-repressed *FLC* increased (Extended Data Fig. 3o-q). Taken together, these data establish that RNAPII-Ser2P is a key determinant of FLD positioning on actively transcribed chromatin.

### Increased gene-body RNAPII-Ser2P and FLD occupancy precedes H3K4me1 erasure

Having established that RNAPII-Ser2P contributes to FLD positioning on active chromatin, we next asked why active transcription recruits a repressive H3K4me1 demethylase without triggering constitutive silencing, and how productive gene-body H3K4me1 erasure is licensed at actively transcribed loci. To address this, we used *FLC* as a model because it exemplifies this paradox: despite being highly transcribed, it retains the capacity to enter stable epigenetic silencing in response to developmental and environmental cues. Moreover, its well-characterized transcriptional and chromatin regulation make it a uniquely tractable system for resolving the sequence of events linking RNAPII dynamics, FLD engagement, and chromatin-state change.

In rapid-cycling Col seedlings, FLD occupancy at silenced *FLC* was minimal, consistent with low RNAPII-Ser2P enrichment across the locus (Extended Data Fig. 4a)^69^. We therefore performed subsequent analyses of FLD chromatin occupancy and H3K4me1 erasure in FRI*-*Col; *pFLD:FLAG-GFP-FLD fld* seedlings, in which the active *FRIGIDA* allele drives high *FLC* expression^84^. In this line, FLD showed substantial chromatin occupancy that was mainly concentrated near the *FLC* TTS, where RNAPII-Ser2P was also prominent, whereas H3K4me1 extended across the gene body but was depleted near the TTS (Fig. 2a)^69,72^. Thus, although FLD is recruited to active *FLC*, the demethylase is spatially offset from much of its gene-body H3K4me1 substrate. Acute inhibition of RNAPII Ser2 phosphorylation with flavopiridol treatment significantly reduced RNAPII-Ser2P and FLD enrichment at the *FLC* TTS (Fig. 2a-c); however, gene-body H3K4me1 was not altered over the same interval at *FLC* (Fig. 2d). A similar pattern was observed genome-wide (Extended Data Fig. 4b,c). These observations indicate that RNAPII-Ser2P-dependent positioning concentrates FLD near TTSs but is not, by itself, sufficient to trigger productive gene-body H3K4me1 erasure.

**Fig. 2.**
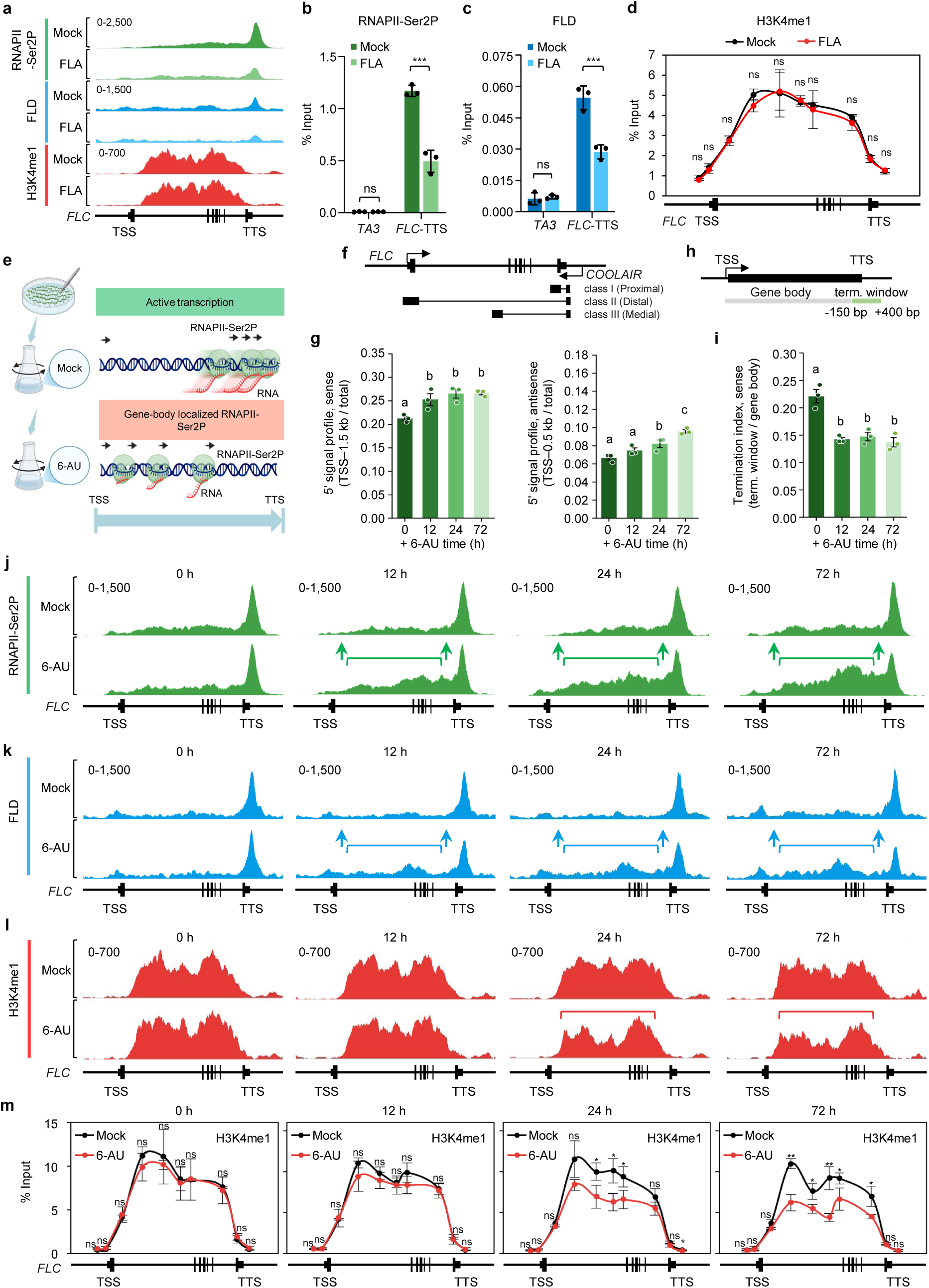
Increased gene-body RNAPII and FLD occupancy precedes H3K4me1 erasure at *FLC*. **a**, IGV tracks of RNAPII-Ser2P, FLD, and H3K4me1 ChIP-seq reads at the *FLC* locus in FRI-Col; *pFLD:FLAG-GFP-FLD fld* seedlings after mock or 1 h flavopiridol (FLA) treatment. The black diagram below shows the *FLC* gene structure. **b**,**c**, ChIP-qPCR validation of RNAPII-Ser2P (**b**) and FLD (**c**) occupancy at the TTS region of the *FLC* locus under the same conditions as in **a**. *TA3* serves as a negative control. Data are mean ± s.d. (*n* = 3 biological replicates). \*\*\**P* < 0.001; ns, not significant, determined by two-tailed Student’s *t*-test. **d**, ChIP-qPCR validation of H3K4me1 levels at the *FLC* locus after flavopiridol treatment. Data are mean ± s.d. (*n* = 3 biological replicates). Statistical significance was determined by two-tailed Student’s *t*-test; ns, not significant. **e**, Schematic illustrating the experimental rationale: the intragenic RNAPII profile during active transcription under mock treatment (top) and the altered RNAPII profile after 6-AU treatment (bottom). **f**, Schematic of the *FLC* locus showing sense transcription and *COOLAIR* antisense transcripts (class I, proximal; class II, distal; class III, medial). **g**, 5′ signal ratio of pNET-seq reads at the *FLC* locus following mock or 6-AU treatment for the indicated times. Left, sense strand, calculated as the fraction of signal within 1.5 kb downstream of the TSS relative to total signal across the *FLC* transcript. Right, antisense strand, calculated as the fraction of signal within 500 bp downstream of the *COOLAIR* TSS relative to total antisense signal. A shorter proximal window was used for the antisense strand owing to the heterogeneity of *COOLAIR* isoform 3′ ends. Data are mean ± s.d. (*n* = 3 biological replicates). Different letters indicate statistically significant differences (*P* < 0.05, one-way ANOVA with Tukey’s post hoc test). **h**, Schematic defining the genomic regions used to calculate the termination index: gene body (bracket) and termination window (−150 bp upstream to +400 bp downstream of the TTS). **i**, Termination index on the *FLC* sense strand, defined as the ratio of pNET-seq reads in the termination window to those in gene-body. Data are mean ± s.d. (n = 3 biological replicates). Different letters indicate statistically significant differences (*P* < 0.05, one-way ANOVA with Tukey’s post hoc test). **j–l**, IGV tracks showing RNAPII-Ser2P (**j**), FLD (**k**) and H3K4me1 (**l**) ChIP-seq singals at the *FLC* locus in FRI-Col; *pFLD:FLAG-GFP-FLD fld* seedlings during the 6-AU time course. Green (**j**) and blue (**k**) arrows indicate increased gene-body occupancy of RNAPII-Ser2P and FLD, respectively; red brackets (**l**) mark the region of gene-body H3K4me1 removal. **m**, ChIP-qPCR validation of H3K4me1 levels at the *FLC* locus under the same conditions as in **j**–**l**. Data are mean ± s.d. (*n* = 3 biological replicates). \**P* < 0.05; ** *P* < 0.01; ns, not significant, determined by two-tailed Student’s *t*-test.

We therefore asked whether productive demethylation depends on a shift in intragenic RNAPII-Ser2P distribution. To test this, we treated FRI*-*Col; *pFLD:FLAG-GFP-FLD fld* seedlings with 6-azauracil (6-AU), which perturbs transcriptional progression by depleting intracellular nucleotide pools^85^, and profiled nascent transcription using pNET-seq alongside ChIP-seq for RNAPII-Ser2P, FLD, and H3K4me1 occupancy throughout a 72-h time course (Fig. 2e). On the sense strand, the 5′ signal ratio (TSS-proximal signal relative to total) increased significantly within 12 h of 6-AU treatment and remained elevated thereafter (Fig. 2f,g). Antisense transcription showed a parallel shift over the same interval (Fig. 2g). Consistently, the termination index, defined as the ratio of read density in the TTS-proximal region to that in the gene body (Fig. 2h), dropped significantly by 12 h and remained low at 24 h and 72 h (Fig. 2i), suggesting a redistribution of elongating RNAPII across the locus.

ChIP-seq analysis revealed a stepwise sequence of chromatin events following this redistribution. After 12 h of 6-AU treatment, RNAPII-Ser2P displayed reduced enrichment at the TTS together with significantly increased occupancy across the gene body (Fig. 2j and Extended Data Fig. 4d,e). FLD chromatin occupancy mirrored RNAPII-Ser2P redistribution (Fig. 2k and Extended Data Fig. 4f), yet gene-body H3K4me1 showed no decrease at this 12-h time point (Fig. 2l,m). By 24 h, RNAPII-Ser2P and FLD remained enriched across the gene body, and this was coincided with the onset of H3K4me1 loss, which became progressively more extensive by 72 h (Fig. 2j-m). Importantly, this elongating RNAPII-dependent H3K4me1 erasure required FLD: in the *fld-4* background, gene-body H3K4me1 at *FLC* did not decrease after 6-AU treatment, consistent with FLD acting as the effector of H3K4me1 removal at *FLC* under these conditions (Extended Data Fig. 4g,h). Thus, perturbation of transcriptional progression is followed first by increased FLD occupancy across the gene body and subsequently by detectable H3K4me1 erasure. Productive H3K4me1 erasure is therefore linked not simply to FLD recruitment to actively transcribed chromatin, but to a redistribution of elongating RNAPII-Ser2P that sustains FLD engagement within the gene body.

Together, these findings suggest that the intragenic occupancy state of elongating RNAPII determines whether FLD remains spatially restricted or gains access to its gene-body H3K4me1 substrate, thereby licensing productive H3K4me1 erasure and the onset of epigenetic silencing.

### The RNAPII-guided chromatin transition recurs during physiological silencing

To determine whether the RNAPII–FLD–H3K4me1 transition observed following chemical perturbation of transcription operates in a physiological silencing context, we examined *FLC* during vernalization, a prolonged cold exposure that represses *FLC* transcription and establishes an epigenetically silenced state maintained after return to warm conditions^86,87^. We profiled *FLC* expression in FRI-Col seedlings and monitored RNAPII-Ser2P, FLD, and H3K4me1 chromatin occupancy in matched FRI*-*Col; *pFLD:FLAG-GFP-FLD fld* plants throughout vernalization. Across all time points examined, *FLD* mRNA and total cellular RNAPII-Ser2P and FLD protein levels remained unchanged (Fig. 3a,b).

**Fig. 3.**
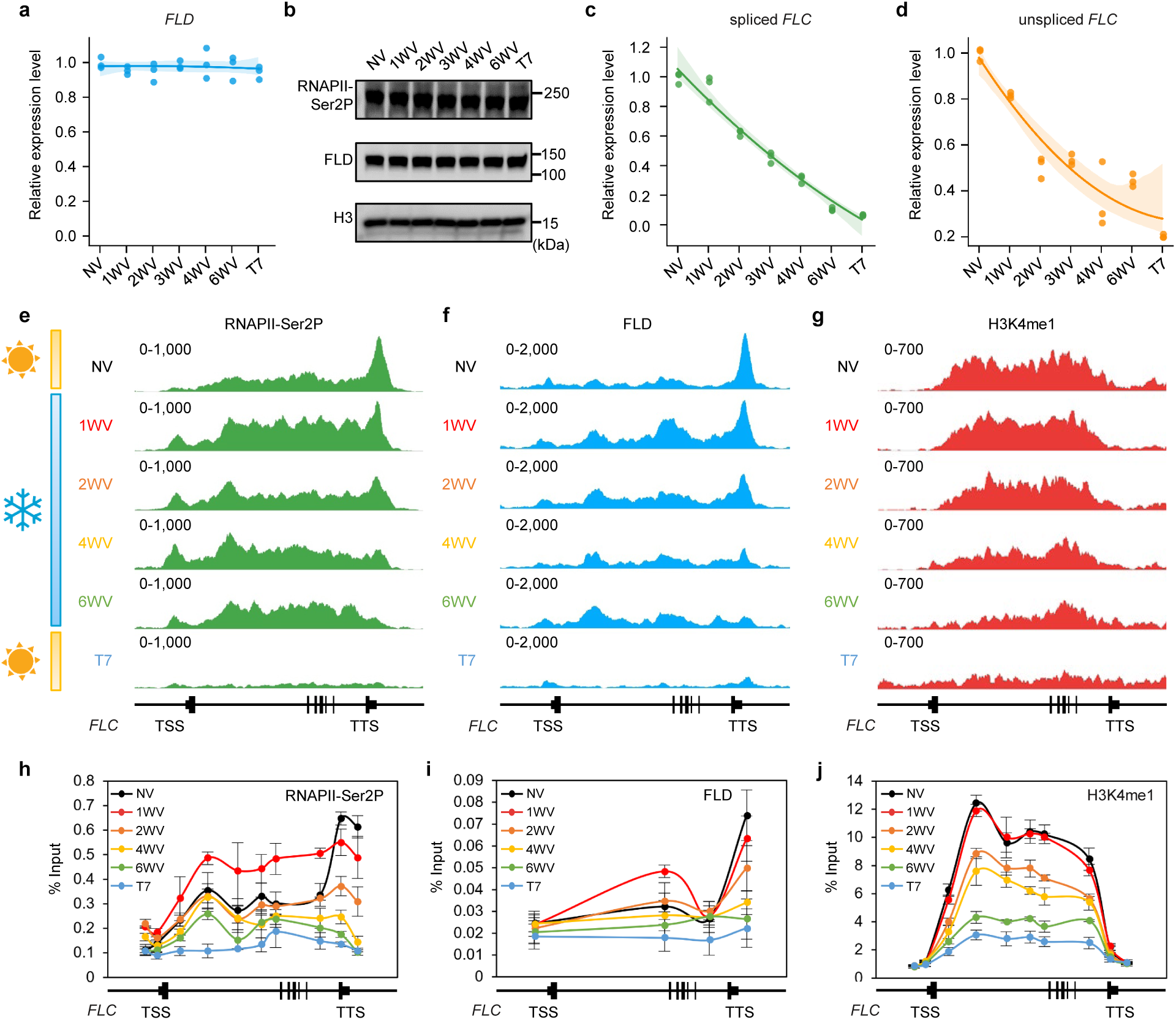
Vernalization increases gene-body RNAPII occupancy to drive FLD engagement and H3K4me1 erasure at *FLC*. **a**, Relative expression of *FLD* measured by RT-qPCR in the FRI-Col background. **b**, Immunoblot showing RNAPII-Ser2P and FLD protein abundance during vernalization (H3, loading control). **c**, **d**, Relative expression of spliced *FLC* (**c**) and unspliced *FLC* (**d**) measured by RT-qPCR in the FRI-Col background. For the RT-qPCR panels (**a**,**c**,**d**): NV, non-vernalized; 1WV–6WV, 1–6 weeks of vernalization; T7, 7 days post-cold; *UBQ10* serves as an internal control; data are mean ± s.d. (*n* = 3 biological replicates). **e**–**g**, ChIP-seq tracks at *FLC* locus for RNAPII-Ser2P (**e**), FLD (**f**) and H3K4me1 (**g**) during vernalization. **h**–**j**, ChIP-qPCR validation of RNAPII-Ser2P (**h**) and FLD (**i**) occupancy and H3K4me1 levels (**j**) at *FLC*. Data are mean ± s.d. (*n* = 3 biological replicates). Gene structure is shown below.

Before cold exposure (NV), *FLC* expression was high (Fig. 3c,d), and the locus showed a 3′-biased RNAPII-Ser2P and FLD occupancy, while H3K4me1 extended across the gene body (Fig. 3e-j). Thus, at transcriptionally active *FLC*, FLD is already recruited yet is spatially separated from much of its gene-body H3K4me1 substrate. After one week of vernalization (1WV), *FLC* expression declined substantially (Fig. 3c,d)^88^. At the 1WV stage, RNAPII-Ser2P ChIP signal increased across the *FLC* gene body relative to NV (Fig. 3e,h), and FLD showed a parallel increase in gene-body occupancy (Fig. 3f,i), whereas gene-body H3K4me1 remained largely unchanged (Fig. 3g,j). As cold exposure continued, gene-body H3K4me1 depletion became detectable until 2 weeks of vernalization (2WV), after RNAPII-Ser2P and FLD had stayed elevated across the gene body for one more week (Fig. 3e-j), placing RNAPII-Ser2P and FLD gene-body relocalization upstream of H3K4me1 erasure during vernalization. Notably, RNAPII-Ser2P and FLD occupancy across the *FLC* gene body started to decrease after 4-6 weeks of cold exposure and eventually disappeared from the chromatin after return to warm conditions, when *FLC* transcription was fully silenced and gene-body H3K4me1 approached near-complete removal (Fig. 3e-j). Altogether, this time-course analysis indicates that RNAPII-Ser2P and FLD relocalize to the gene body before detectable H3K4me1 erasure during vernalization, establishing FLD gene-body redistribution as an early event in epigenetic silencing. Once the gene becomes silenced, FLD disengages from chromatin, indicating that active transcription is required for its continued chromatin association.

We next asked whether RNAPII-Ser2P-dependent gene-body FLD engagement and H3K4me1 erasure is specific to vernalization-induced silencing or instead reflects a broader requirement during silencing transitions. In the rapid-cycling Col accession, *FLC* expression in dry seeds was very high but declined sharply within 48 h of imbibition and remained silenced in seedlings (Fig. 4a). The decline in *FLC* expression was largely attenuated in *fld* mutants, and this defect was accompanied by the persistence of gene-body H3K4me1 relative to Col (Fig. 4a,b). Thus, FLD is required for gene-body H3K4me1 erasure at *FLC* during this silencing transition. To determine when FLD is required, we used an estradiol-inducible *aMIRFLD* line that enables acute *FLD* depletion at defined stages of the silencing process, allowing us to distinguish its requirement for silencing initiation from that for silencing maintenance (Fig. 4c). Treating dry seeds with estradiol reduced *FLD* mRNA within 48 h of germination and slowed the early decline in *FLC* expression (Fig. 4d-f). By contrast, although induction in 5-day-old seedlings similarly reduced *FLD* mRNA, when *FLC* was already silenced, FLD depletion for up to four days did not reactivate *FLC* (Fig. 4g-i). Thus, FLD is required for initiating gene silencing during the early high-to-low transcriptional transition but becomes dispensable once the transcriptionally inactive state has been established.

**Fig. 4.**
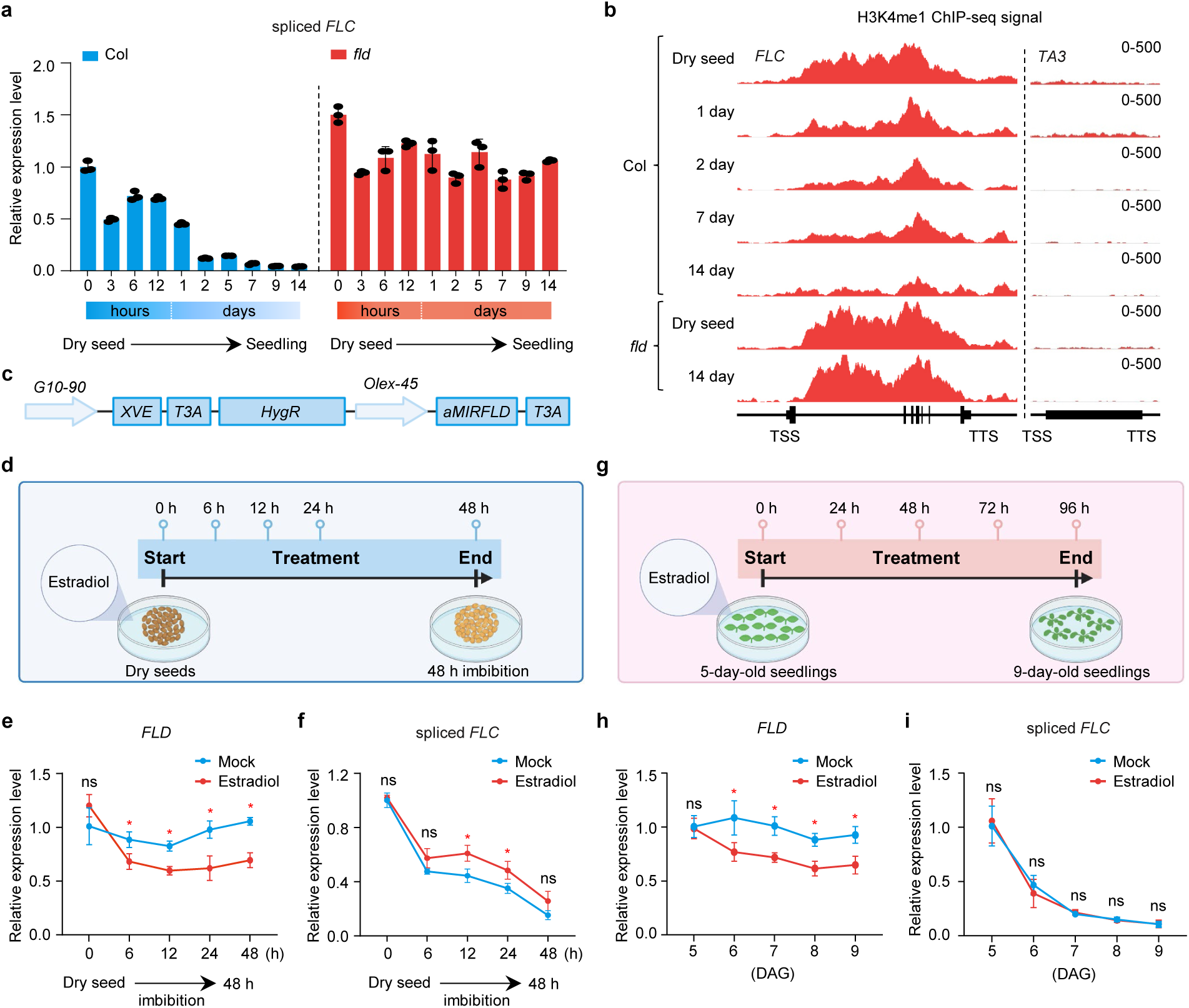
FLD promotes post-germination *FLC* repression in rapid-cycling Col through gene-body H3K4me1 removal. **a**, Relative expression of spliced *FLC* in Col (left) and *fld-4* (right) from dry seeds to 14-day-old seedlings measured by RT-qPCR. Dashed line separates genotypes. The x axis shows time post-imbibition in hours (0, 3, 6, and 12 h) followed by days (1, 2, 5, 7, 9, and 14 d). Data are mean ± s.d. (*n* = 3 biological replicates). **b**, H3K4me1 ChIP-seq tracks at *FLC* and *TA3* (negative control) in Col across post-germination development (dry seed, and 1-, 2-, 7- and 14-day-old seedlings) and in *fld-4* at the dry-seed and 14-day-old seedlings. **c**, Schematic of the estradiol-inducible *aMIRFLD* knockdown construct. **d**,**g**, Experimental design for estradiol-induced FLD knockdown in dry seeds (**d**) and 5-day-old seedlings (**g**). **e**,**f**, Relative expression of *FLD* (**e**) and spliced *FLC* (**f**) following estradiol treatment of dry seeds during 48 h of imbibition. Mock serves as a vehicle control. Data are mean ± s.d. (*n* = 3 biological replicates). Statistical significance was determined by two-tailed Student’s *t*-test; ns, not significant; \**P* < 0.05. **h**,**i**, Relative expression of *FLD* (**h**) and spliced *FLC* (**i**) following estradiol treatment of 5-day-old seedlings monitored to 9 days after germination (DAG). Data are mean ± s.d. (*n* = 3 biological replicates). Statistical significance was determined by two-tailed Student’s *t*-test; ns, not significant; \**P* < 0.05.

Together, these analyses suggest a common physiological silencing transition in which redistribution of RNAPII-Ser2P precedes FLD-dependent gene-body H3K4me1 erasure. Importantly, FLD functions during silencing initiation but becomes dispensable once stable repression has been achieved, indicating that the RNAPII– FLD–H3K4me1 chromatin transition acts during silencing entry rather than maintenance.

### FLD engages RNAPII-Ser2P on chromatin through LD

Identification of RNAPII-Ser2P-mediated positioning of the silencing-promoting FLD demethylase to convert transcription into gene-body H3K4me1 removal for gene silencing raised the question of how FLD is physically linked to elongating RNAPII-Ser2P on chromatin. FLD was recently reported to form a multiprotein complex with LUMINIDEPENDENS (LD), SET DOMAIN GROUP 26 (SDG26), EARLY FLOWERING 4-like 2/4 (EFL2/4), and ANTHESIS PROMOTING FACTOR 1 (APRF1)^71,72,89^. Structural modelling using AlphaFold3 predicted that FLD directly interacted with LD, but not with the other subunits (Fig. 5a), consistent with recent work^89^. Given its direct interaction with FLD, we reasoned that LD might be required for FLD chromatin engagement. To test this, we introduced the *ld* mutation into the *pFLD:FLAG-GFP-FLD* transgenic line. Consistent with a previous report^71^, loss of LD reduced FLD protein abundance, and we further found that it caused a near-complete loss of FLD chromatin occupancy (Fig. 5b,c and Extended Data Fig. 5a-c). Proteasome inhibition with MG132 restored FLD protein levels in the *ld* background (Fig. 5b), but failed to rescue FLD chromatin association: FLD occupancy remained severely reduced at more than 10,000 FLD target loci and was nearly completely lost at TTSs of RNAPII-Ser2P-enriched convergent genes (Fig. 5d-g and Extended Data Fig. 5b,c and Supplementary Table 2). These results demonstrate that LD is required for FLD chromatin association, independently of its role in maintaining FLD protein stability. We also evaluated whether LD chromatin occupancy reciprocally depends on FLD. Genome-wide LD occupancy in *fld* mutant background was indistinguishable from that in wild-type (Fig. 5h,i and Extended Data Fig. 5d,e). Notably, at *FLC*, where transcription is de-repressed in *fld*, LD occupancy increased (Fig. 5j-m), consistent with transcription-coupled, rather than FLD-dependent, LD chromatin association.

**Fig. 5.**
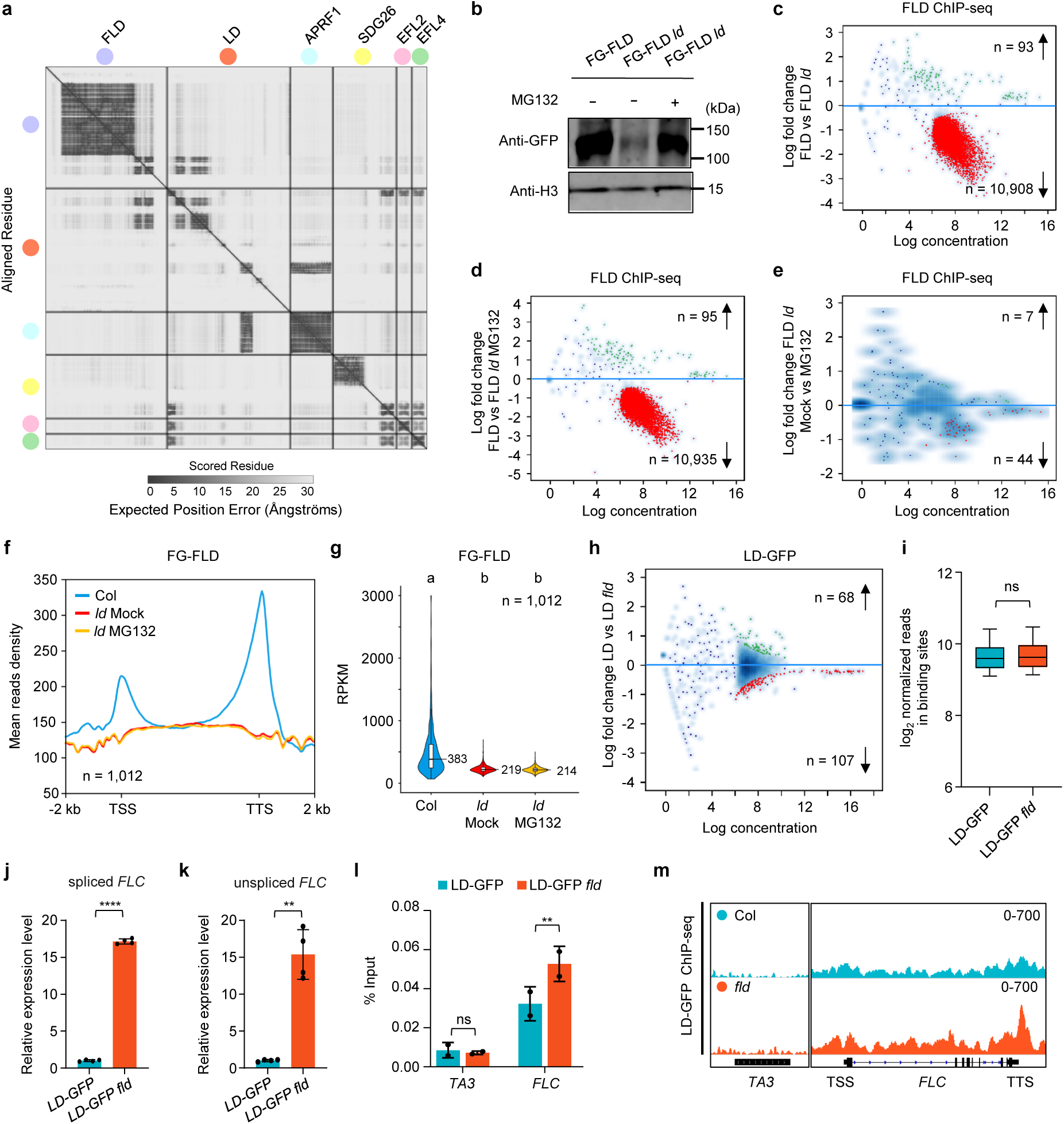
FLD engages RNAPII-Ser2P on chromatin through LD. **a**, Predicted aligned error (PAE) plot from an AlphaFold 3 prediction of the full FLD complex, shown as a reference. Subunits are colored as indicated: FLD (purple), LD (red), APRF1 (cyan), SDG26 (yellow), EFL2 (pink) and EFL4 (green). **b**, Immunoblot showing FLD protein levels in FG-FLD, FG-FLD *ld* and FG-FLD *ld* treated with MG132. H3 serves as a loading control. **c**–**e**, Scatter plots of FLD ChIP-seq occupancy. **c**, FLD in Col versus *ld* (*n* = 10,908 decreased, *n* = 93 increased; FDR < 0.05). **d**, FLD in Col versus *ld* + MG132 (*n* = 10,935 decreased, *n* = 95 increased). **e**, FLD in *ld* mock versus *ld* + MG132 (*n* = 44 decreased, *n* = 7 increased). **f**,**g**, Metagene plot (**f**) and violin plot (**g**) of FLD ChIP-seq signal at RNAPII-Ser2P target genes meeting the criteria in Fig. 1d (*n* = 1,012) in the indicated backgrounds. Different letters indicate significant differences (one-way ANOVA with Tukey’s post hoc test). **h**, Scatter plot of LD-GFP ChIP-seq occupancy in Col versus *fld*. Red, decreased; green, increased (FDR < 0.05). **i**, Box plot of LD ChIP-seq read density at binding sites. ns, not significant, determined by two-tailed Student’s *t*-test. **j**,**k**, Relative expression of spliced *FLC* (**j**) and unspliced *FLC* (**k**) in LD-GFP lines in Col and *fld* backgrounds at 7 DAG. Data are mean ± s.d. (*n* = 4 biological replicates). Statistical significance was determined by two-tailed Student’s *t*-test; \*\**P* < 0.01; \*\*\*\**P* < 0.0001. **l**, ChIP-qPCR validation of LD occupancy at the indicated loci in Col and *fld* backgrounds. *TA3* serves as a negative control. Data are mean ± s.d. (*n* = 2 biological replicates). Statistical significance was determined by two-tailed Student’s *t*-test; \*\**P* < 0.01; ns, not significant. **m**, LD ChIP-seq tracks at *TA3* (negative control) and *FLC* in Col and *fld* backgrounds.

Together, these data identify LD as the subunit required for FLD chromatin association, whereas LD itself associates with chromatin independently of FLD.

### The C-terminal region of FLD is required for FLD chromatin engagement through direct interaction with LD

To dissect how LD promotes FLD chromatin occupancy, we further examined AlphaFold-guided structural models of the FLD–LD complex. The predicted model suggested that FLD interacts with LD through a C-terminal region of unknown function (hereafter named FLD-C) (Figs. 5a and 6a), consistent with a previous report^89^. Notably, FLD is distinguished from other LSD1 orthologs and its Arabidopsis paralogs LDL1/2/3 by this unique C-terminal region, which is specific to land plants and conserved across the land plant species examined here (Fig. 6b and Extended Data Fig. 6a,b). *In vitro* pull-down assays confirmed that FLD-C directly interacted with LD (Extended Data Fig. 6c).

**Fig. 6.**
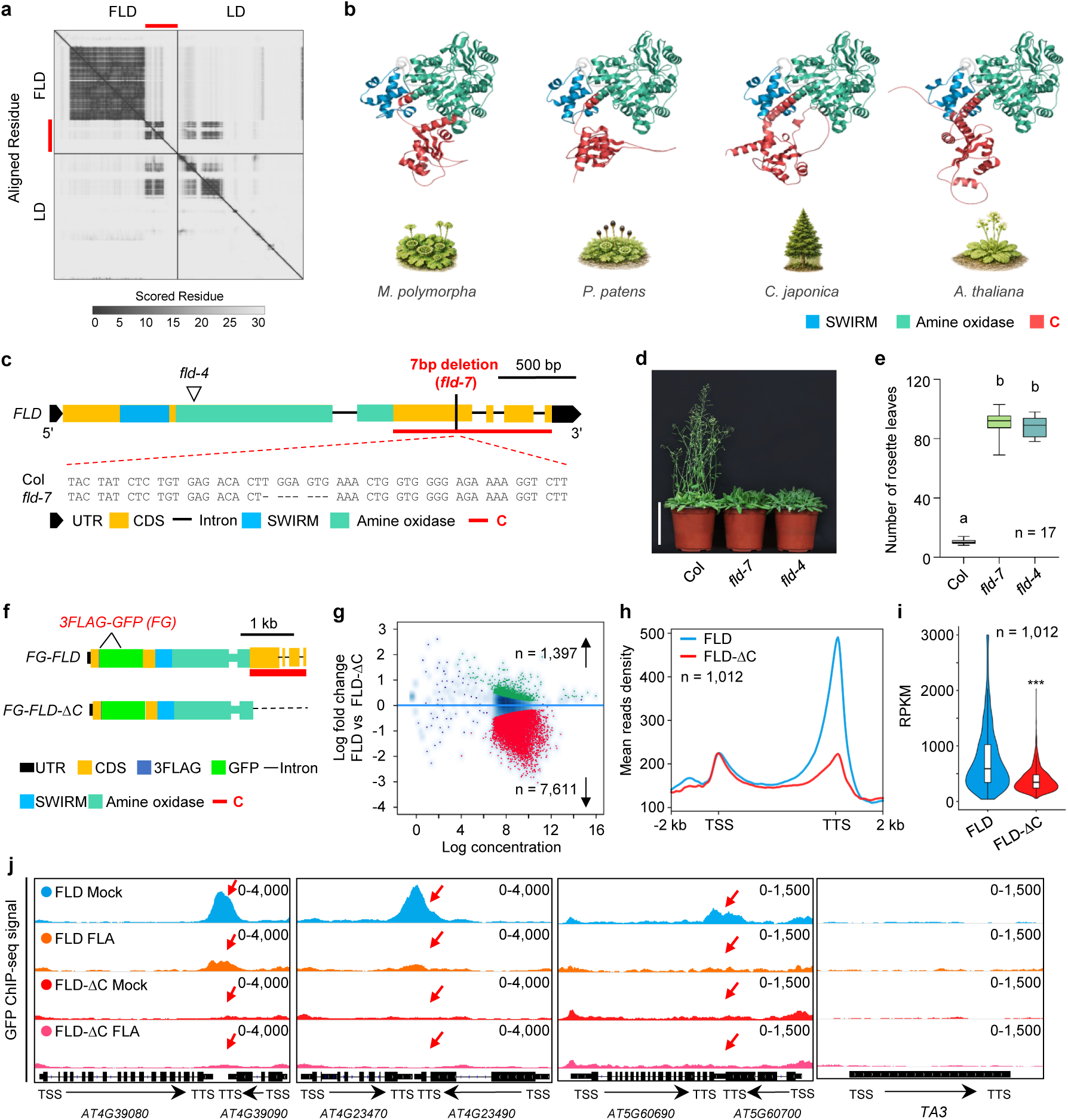
The C-terminal region of FLD is required for FLD chromatin engagement through direct interaction with LD. **a**, PAE plot from an AlphaFold 3 prediction of the direct interaction between FLD and LD. Red line indicates FLD C-terminal that interacts with LD. **b**, AlphaFold-predicted structures of FLD homologues from *Marchantia polymorpha* (liverwort), *Physcomitrium patens* (moss), *Cryptomeria japonica* (gymnosperm) and *Arabidopsis thaliana* (angiosperm). The SWIRM (blue) and amine oxidase (green) domains are shown, and the C-terminal region (red) becomes progressively structured across the examined land plants from bryophytes to angiosperms. **c**, Schematic of *FLD* gene structure. Black triangle, T-DNA insertion site (*fld-4*); black line, 7-bp deletion (*fld-7*). Known domains (SWIRM and amine oxidase) and the plant-specific C-terminal region (red) are indicated. Bottom, sequence alignment of *fld-7* versus Col. **d**, Flowering phenotypes of Col and *fld* mutants grown under long-day conditions for 5 weeks. Scale bar, 10 cm. **e**, Rosette leaf number at bolting. Different letters indicate significant differences (one-way ANOVA with Tukey’s post hoc test). *n* = 17 plants. **f**, Schematic of the transgene constructs FG-FLD (full-length) and FG-FLD-ΔC (C-terminal deletion). **g**, Scatter plot of FLD versus FLD-ΔC ChIP-seq occupancy. Differentially occupied peaks with a false discovery rate (FDR) < 0.05 are highlighted in green (increased, *n* = 1,397) and red (decreased, *n* = 7,611). **h**, Metagene plot of FLD and FLD-ΔC ChIP-seq signals at RNAPII-Ser2P target genes meeting the criteria in Fig. 1D (*n* = 1,012). **i**, Violin plot of ChIP-seq read density at the same genes. \*\*\**P* < 0.001 (two-tailed Student’s *t*-test). **j**, ChIP-seq tracks of FG-FLD and FG-FLD-ΔC at representative RNAPII-Ser2P target genes after mock or flavopiridol treatment. Red arrows indicate the positions of FLD peaks.

To examine the function of FLD-C *in vivo*, we generated a *fld* mutant allele (*fld-7*) carrying a 7-bp deletion that causes a frameshift and premature termination, resulting in truncation of FLD-C (Fig. 6c and Extended Data Fig. 7a-c). *fld-7* phenocopied the loss-of-function allele *fld-4*, showing extremely delayed flowering and increased *FLC* expression (Fig. 6d,e and Extended Data Fig.7d,e). *fld-7* also resembled *fld-4* at the chromatin level, causing genome-wide increases in H3K4me1, including at *FLC* (Extended Data Fig. 7f-i). Full-length FLD transgene complemented the *fld-7* phenotypes (Extended Data Fig. 8a-e). These results indicate that disruption of the C-terminal region abolishes FLD function *in vivo*.

To test whether FLD-C is required for FLD chromatin association, we expressed a truncated version of FLD lacking the C-terminal region (FLD-ΔC; amino acids 672– 936 deleted) in the *fld-7* background (Fig. 6f). In contrast to full-length FLD, FLD-ΔC failed to restore flowering time, *FLC* repression, and H3K4me1 demethylation (Extended Data Fig. 8a-f). FLD-ΔC protein accumulated to levels comparable to those of full-length FLD (Extended Data Fig. 8g-j); however, its occupancy was reduced at 7,611 FLD target sites, far exceeding the 1,397 sites at which FLD-ΔC occupancy increased (Fig. 6g and Extended Data Fig. 8k,l). Importantly, at convergently transcribed genes, loss of the C terminus severely reduced FLD enrichment near transcription termination sites (Fig. 6h-i), indicating that FLD engagement at termination-proximal regions requires FLD-C. We also explored whether this requirement acts within RNAPII-Ser2P-dependent FLD chromatin positioning. Flavopiridol treatment reduced cellular RNAPII-Ser2P levels without altering FLD or FLD-ΔC protein abundance (Extended Data Fig. 9a) and lowered full-length FLD chromatin occupancy (Extended Data Fig. 9b-d). By contrast, acute reduction of RNAPII Ser2 phosphorylation did not further reduce FLD-ΔC chromatin occupancy (Fig. 6j and Extended Data Fig. 9d-f). Together, these results indicate that FLD-C is required for RNAPII-Ser2P-dependent chromatin engagement.

In summary, these data demonstrate that FLD engages RNAPII-Ser2P-enriched chromatin through an interaction between its distinctive C-terminal region and LD.

### The LD TFIIS domain mediates Ser2P-sensitive coupling of the FLD complex to elongating RNAPII

We next asked how LD is linked to RNAPII-Ser2P. AlphaFold-based structural modelling predicted a direct interaction between the C-terminal region of LD and APRF1 (Fig. 5a), consistent with previous findings that, together with TOPP4, these proteins form a CPF-like phosphatase complex^72^. Based on this interaction, LD was proposed to promote *FLC* silencing through CPF^72^. However, *FLC* silencing is more severely affected by loss of LD than by loss of CPF^72^ or deletion of the LD C-terminal region (Fig. 7a-e and Extended Data Fig. 10a,b). These observations suggest that a model involving CPF only partially explains the function of LD and that LD might exert its key role in gene silencing also through other, CPF-independent pathways. In addition to the C-terminal region that interacts with APRF1, LD contains an N-terminal homeodomain (HOX) and a central TFIIS domain^72,89^. We therefore examined how LD might be recruited to chromatin by testing whether these two domains mediate its targeting to chromatin. We stably expressed truncated versions of LD lacking the HOX or the TFIIS domain from the native *LD* promoter in *ld* mutant plants (*pLD:LD-ΔHOX-GFP ld* and *pLD:LD-ΔTFIIS-GFP ld*) (Fig. 7a and Extended Data Fig.10a). We found that both the HOX-deleted and TFIIS-deleted versions of LD failed to silence *FLC*, with plants remaining late flowering, similar to the *ld* loss-of-function mutant (Fig.7b-e and Extended Data Fig. 10b), suggesting that both domains are essential for the biological function of LD.

**Fig. 7.**
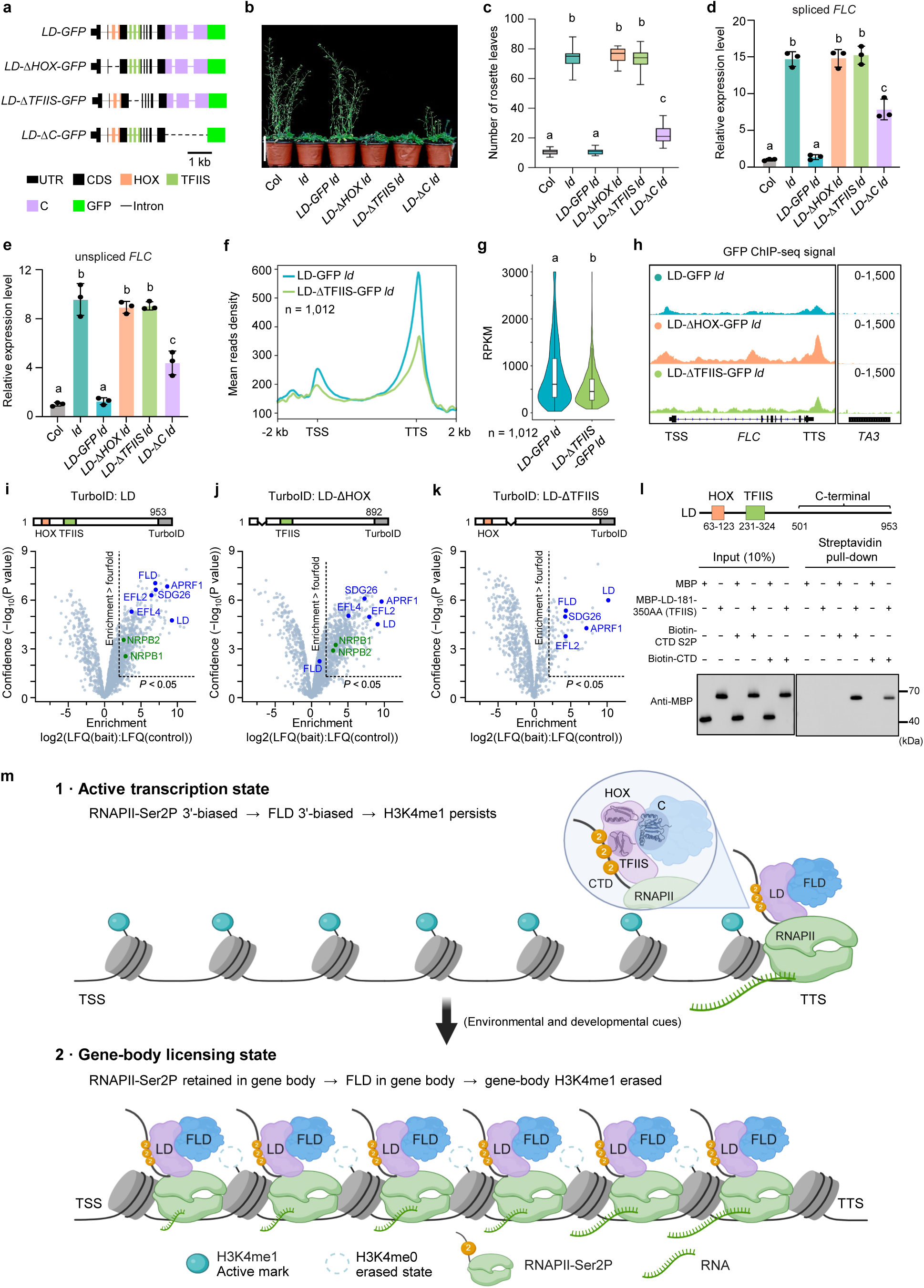
The LD TFIIS domain mediates Ser2P-sensitive coupling of the FLD complex to elongating RNAPII. **a**, Schematic of GFP-tagged LD constructs: full-length LD-GFP, LD-ΔHOX-GFP (HOX domain deletion), LD-ΔTFIIS-GFP (TFIIS domain deletion), and LD-ΔC-GFP (C-terminal deletion). **b**, Flowering phenotypes of Col, *ld*, *LD-GFP ld*, *LD-ΔHOX-GFP ld*, *LD-ΔTFIIS-GFP ld*, and *LD-ΔC-GFP ld* plants grown under long-day (LD) conditions for 5 weeks. Scale bar, 10 cm. **c**, Rosette leaf number at bolting. Different letters indicate significant differences (one-way ANOVA with Tukey’s post hoc test). **d**,**e**, Relative expression of spliced (**d**) and unspliced (**e**) *FLC* in the indicated genotypes at 7 DAG, measured by RT-qPCR. Data are mean ± s.d. (*n* = 3 biological replicates). Different letters indicate significant differences (one-way ANOVA with Tukey’s post hoc test). **f**, **g**, Metagene plot (**f**) and violin plot (**g**) of LD-GFP and LD-ΔTFIIS-GFP ChIP-seq signals at RNAPII-Ser2P target genes meeting the criteria in Fig. 1d (*n* = 1,012) in the *ld* background. Different letters indicate significant differences. **h**, ChIP-seq tracks at *FLC* and *TA3* for LD-GFP, LD-ΔHOX-GFP, and LD-ΔTFIIS-GFP in the *ld* background. **i**–**k**, Volcano plots of TurboID-based proximity labelling for full-length LD (**i**), LD-ΔHOX (**j**) and LD-ΔTFIIS (**k**), each compared with the *p35S:GFP-TurboID* non-specific labelling control. Above each plot, the corresponding TurboID fusion construct is shown. FLD–LD complex subunits (blue) and RNAPII subunits (green) are labelled. Dashed lines indicate enrichment > fourfold and *P* < 0.05. **l**, Streptavidin pull-down assay testing the interaction between the LD TFIIS domain (amino acids 181–350) and Ser2-phosphorylated (biotin-CTD-S2P) or unphosphorylated (biotin-CTD) CTD heptapeptides. Above, schematic of LD protein domain architecture. **m**, Model for a gene-body RNAPII-Ser2P occupancy checkpoint coupled to FLD–LD-mediated H3K4me1 erasure. **1, Active transcription state.** At actively transcribed loci, RNAPII-Ser2P occupancy and FLD–LD engagement is biased toward the 3′/TTS-proximal region, whereas H3K4me1 spans the gene body. Under this configuration, FLD is present on chromatin, but limited gene-body RNAPII-Ser2P occupancy restricts productive FLD–LD engagement across the H3K4me1-marked substrate, thereby constraining H3K4me1 erasure. **2, Gene-body licensing state.** Transcriptional perturbation or physiological silencing cues induces redistribution of RNAPII-Ser2P into the gene body, potentially integrating sense and antisense transcriptional inputs where present. Increased gene-body RNAPII-Ser2P sustains productive FLD–LD engagement over the H3K4me1-marked nucleosomes, enabling FLD-dependent erasure of gene-body H3K4me1. The FLD–LD complex functions as a modular system that separates RNAPII coupling, FLD–LD assembly, and catalytic activity. The LD TFIIS domain engages the RNAPII CTD in a Ser2P-sensitive manner, coupling the complex to elongating RNAPII and supporting LD chromatin occupancy. FLD-C docks FLD onto LD, whereas the FLD catalytic core, comprising the SWIRM and amine oxidase domains, removes gene-body H3K4me1.

We then compared the chromatin occupancy of LD-ΔHOX and LD-ΔTFIIS with that of full-length LD. Compared to full-length LD, LD-ΔTFIIS showed significantly reduced occupancy at more than 7,000 LD-bound sites (Extended Data Fig. 10c and Supplementary Table 3). This reduction was observed at transcription termination sites where RNAPII-Ser2P enriched (Fig. 7f,g). In contrast, LD-ΔHOX-GFP showed genome-wide occupancy similar to that of full-length LD-GFP (Extended Data Fig. 10d,e), suggesting that HOX is dispensable for LD chromatin occupancy. These data indicate that the TFIIS but not the HOX domain of LD mediates LD targeting to RNAPII-Ser2P-enriched chromatin.

We further examined the contribution of the TFIIS domain to LD occupancy at *FLC*. As noted above, the two truncation lines highly expressed *FLC* at comparable levels such that they provide a matched transcriptional context in which differences in LD occupancy do not simply reflect differences in transcription status (Fig. 7d). Under these conditions, full-length LD-GFP showed minimal enrichment at the silenced *FLC*, whereas LD-ΔHOX was enriched at transcriptionally active *FLC*, particularly at the TTS where RNAPII-Ser2P accumulates (Fig. 7h and Extended Data Fig. 10f). By contrast, LD-ΔTFIIS chromatin occupancy was minimal, even though *FLC* was similarly transcriptionally reactivated (Fig. 7h and Extended Data Fig. 10f). These data further support a selective requirement for the TFIIS domain in LD chromatin targeting. To examine whether the TFIIS domain could mediate LD’s engagement with RNAPII-Ser2P, we performed comparative TurboID-based proximity labelling using full-length LD and LD truncation variants lacking either the HOX or TFIIS domain (Fig. 7i-k). Full-length LD robustly enriched all known components of the FLD–LD complex, including FLD, APRF1, SDG26 and EFL2/4, as well as the two largest RNAPII subunits, NRPB1 and NRPB2 (Fig. 7i), consistent with LD coupling the FLD complex to RNAPII. Deletion of the HOX domain did not significantly affect the enrichment of NRPB1, NRPB2, APRF1, SDG26 or EFL2/4, but specifically and markedly reduced FLD enrichment (Fig. 7j), suggesting that the HOX domain mediates the interaction between LD and FLD. In contrast, deletion of the TFIIS domain preserved enrichment of FLD and other FLD–LD complex components, including APRF1, SDG26 and EFL2, but abrogated enrichment of NRPB1 and NRPB2 (Fig. 7k), indicating that association of LD with RNAPII depends on the TFIIS domain. We next tested whether the TFIIS domain could directly bind RNAPII CTD. In peptide pull-down assays, recombinant LD-TFIIS bound both unmodified and Ser2-phosphorylated CTD peptides, with consistently stronger recovery observed for the Ser2-phosphorylated peptide (Fig. 7l), suggesting that LD TFIIS domain can directly bind to the RNAPII CTD and that this binding is enhanced by Ser2 phosphorylation.

Taken together, these data strongly suggest that the LD TFIIS domain directly engages the RNAPII CTD in a Ser2P-sensitive manner, thereby coupling the FLD–LD complex to elongating RNAPII on chromatin, whereas the HOX domain mediates LD association with FLD.

## Discussion

Transcription is generally viewed as a mechanism that reinforces active chromatin states. Yet developmental and cell fate transitions frequently require actively transcribed loci to enter stable epigenetic silencing. Our findings reveal an unexpected solution to this paradox. We show that a shift in the intragenic distribution of RNAPII can be coupled to a repressive chromatin effector, the FLD–LD H3K4me1 demethylase complex, during entry into a silencing-competent state. Critically, the complex does not simply track polymerase presence but interprets its intragenic distribution, such that productive H3K4me1 removal is licensed when RNAPII exhibits sustained occupancy across gene bodies. These results extend the conceptual scope of co-transcriptional chromatin regulation from maintaining active chromatin states to governing their dismantling. More broadly, they identify intragenic RNAPII occupancy states as potential decision signals that determine whether an actively transcribed locus remains active or becomes competent for epigenetic silencing.

Mechanistically, our data address a fundamental question of how an actively transcribed locus can recruit a repressive H3K4 demethylase yet avoid constitutive silencing. RNAPII-Ser2P is required for FLD chromatin association (Fig. 1f-n). However, recruitment alone is insufficient for productive demethylation. Under high transcription conditions, RNAPII-Ser2P concentrates FLD occupancy near transcription termination sites, whereas H3K4me1 occupies the gene body (Fig. 2a). Productive demethylation is therefore constrained by the spatial separation between FLD enzyme and its H3K4me1 substrate. Following transcriptional perturbation, increased gene-body RNAPII-Ser2P occupancy—as observed after 6-AU treatment (Fig. 2j) or during early vernalization (Fig. 3e,h)—is accompanied by a parallel redistribution of FLD (Figs. 2k and 3f,i), whereas gene-body H3K4me1 decline occurs only subsequently (Figs. 2l,m and 3g,j). This temporal sequence supports an RNAPII-distribution checkpoint model in which RNAPII occupancy patterns gate access of a chromatin eraser to its substrate, thereby controlling entry into a silencing-permissive chromatin state.

Co-transcriptional chromatin modification is well established as a mechanism for reinforcing active transcription, exemplified by RNAPII-coupled H3K4^11-17^ and H3K36 methylation pathways^18-24^. In budding yeast, RNAPII-coupled H3K36 methylation recruits the Rpd3S histone deacetylase complex to transcribed regions, where it suppresses spurious intragenic transcription and preserves transcriptional fidelity rather than initiating stable epigenetic silencing^23,24^. The *Arabidopsis* H3K4me2 demethylase LDL3 likewise exemplifies the conventional mode of co-transcriptional chromatin regulation, in which LDL3 associates with phosphorylated RNAPII to support active transcription^90^. By contrast, the mechanism described here couples ongoing transcription to achieve the opposite chromatin outcome. This reversal is enabled by a modular architecture that separates polymerase coupling, FLD–LD assembly, and catalytic activity, with LD functioning as an adaptor that couples the FLD demethylase to elongating RNAPII (Figs. 5 and 7l,m). The FLD–LD system therefore repurposes RNAPII coupling from reinforcing active chromatin to promoting its dismantling during entry into epigenetic silencing.

Previous studies have established important roles for antisense transcription and co-transcriptional RNA processing during *FLC* silencing^71,72,81,88,91^. Our findings identify an unprecedented regulatory layer centered on elongating RNAPII-Ser2P. Whereas previous models emphasized information conveyed by RNA molecules and their processing, our results demonstrate that the intragenic positioning of elongating RNAPII-Ser2P can also provide an instructive input into chromatin-state transitions. Because FLD–LD is coupled to RNAPII through CTD recognition rather than transcriptional direction, sense and antisense transcription may jointly shape the RNAPII landscape read by the FLD complex. Notably, LD also associates with multiple RNA processing factors, including FCA and FPA^71,72,81^ (Supplementary Table 3), placing LD at the molecular interface between RNAPII-Ser2P-dependent FLD positioning and co-transcriptional RNA processing. How these regulatory inputs are mechanistically coordinated remains to be determined.

Importantly, analyses of both vernalization and post-germination downregulation indicate that this RNAPII-distribution checkpoint is reused across distinct physiological settings. During vernalization, gene-body RNAPII-Ser2P–FLD engagement precedes detectable gene-body H3K4me1 erasure (Fig. 3e-j). Likewise, during post-germination downregulation, FLD is required for the high-to-low transcriptional transition but is largely dispensable once *FLC* silencing has been established (Fig. 4f-i). Together, these observations suggest that distinct environmental and developmental inputs converge on a common chromatin-entry route and that RNAPII occupancy states represent a general regulatory layer through which diverse upstream signals are integrated before stable epigenetic transitions occur. This places the polymerase-distribution checkpoint within the analog-to-digital framework proposed for *FLC* silencing^72,87,92-96^: dynamic changes in RNAPII occupancy and distribution regulate entry into a silencing-permissive state, whereas Polycomb mechanisms can subsequently stabilize the epigenetically OFF state. Identifying the mechanisms that reshape intragenic RNAPII occupancy states should therefore further reveal how transcriptional states are converted into stable chromatin-state changes during development and environmental adaptation.

Our temporal dissection at *FLC*, together with genome-wide analyses of RNAPII-Ser2P, FLD, and LD occupancy, suggests that the RNAPII-dependent silencing mechanism operates broadly across the genome. In line with this, recent work identified LD as a regulator of genome-wide transcriptional repression^97^. Together, these findings suggest that RNAPII-dependent positioning of the FLD–LD complex represents a broadly used mechanism for coupling ongoing transcription to epigenetic silencing and raise the question of what determines local RNAPII occupancy states at individual loci. Because the checkpoint appears to respond to local RNAPII occupancy rather than bulk elongation rate, *cis*-encoded features such as nascent RNA structure and R-loop-prone regions may help define the RNAPII landscapes interpreted by the FLD–LD complex^98-101^. More broadly, this framework now makes it possible to ask how cis-encoded features shape the probability of epigenetic switching at developmentally regulated genes.

In summary, our findings reveal a paradoxical mode of co-transcriptional chromatin regulation in which an elongating polymerase guides a silencing-promoting epigenetic eraser to remove the very active chromatin state that transcription helps sustain. Rather than functioning solely as a readout of gene activation, elongating RNAPII can act as an instructive platform whose occupancy state helps determine when and where chromatin resetting occurs. We propose that intragenic RNAPII occupancy states function as chromatin-state decision signals that couple ongoing transcription to the initiation of epigenetic silencing. Through this mechanism, active transcription becomes not only a hallmark of the active state but also a licensing mechanism for its epigenetic shutdown.

## Materials and Methods

### Plant materials and growth conditions

The *Arabidopsis thaliana* Columbia-0 (Col-0, hereafter Col) ecotype was used as the wild-type background for all experiments. The FRI-Col line containing an active *FRIGIDA* allele introgressed into the Col background was used for vernalization and *FLC* expression studies^84^. Col lacks a functional *FRIGIDA* allele and was used where indicated as a rapid-cycling background with low basal *FLC* expression. Experiments requiring high *FLC* expression, including the analyses shown in Figs. 2 and 3, were performed in the FRI-Col background or in FRI-Col-derived transgenic or mutant lines unless otherwise stated.

Loss-of-function mutants used in this study include *fld-4* (SALK_015053) and *ld* (SALK_003039)^89^. The CRISPR-induced *fld-7* allele carries a 7-bp deletion within the region encoding the C terminus, causing a translational frameshift from amino acid position 801 onward and a premature stop codon shortly thereafter. Amino acid numbering refers to the FLD.3 isoform (936 aa; AT3G10390.3). Seeds were surface sterilized, sown onto half-strength Murashige and Skoog (1/2 MS) medium containing 1% sucrose and 0.8% agar, and stratified at 4 °C in the dark for 3 days before transfer to controlled-environment chambers under long-day conditions (16 h light/8 h dark) at 22°C. Homozygous T-DNA insertion mutants were identified by PCR-based genotyping.

### Vernalization treatment

Seeds were surface sterilized and sown onto 1/2 MS medium plates, then stratified at 4°C in the dark for 3 days. For non-vernalized (NV) conditions, seedlings were grown for 7 days under long-day conditions (16 h light/8 h dark, constant 22°C) before sample collection. For vernalization treatment, seedlings were first pre-grown for 7 days under long-day conditions (16 h light/8 h dark, constant 22°C) and then transferred to vernalization conditions at 4°C under short-day photoperiods (8 h light/16 h dark) for the indicated durations (1, 2, 4, or 6 weeks). For transfer experiments (T7), after 6 weeks of cold treatment, seedlings were returned to long-day conditions (16 h light/8 h dark, constant 22°C) on the plates for 7 days of recovery. Samples were collected at the specified time points for subsequent analyses.

### Generation of transgenic plants

For the *pFLD:FLAG-GFP-FLD* construct, the full-length *FLD* genomic sequence including a 2,699-bp native promoter (upstream of the ATG start codon) and the coding region was fused with FLAG-GFP and cloned into the *pEarleyGate301* destination vector through the Gateway cloning system to generate an N-terminal FLAG-GFP fusion. For truncation constructs, FLD-ΔC lacks the C-terminal region (amino acids 672–936 deleted). The *fca-12* allele was generated by CRISPR/Cas9 in the Col background and carries a 1-bp insertion in *FCA* (AT4G16280) that causes a frameshift and a premature stop codon (Extended Data Fig. 2a). *fpa-12* is a T-DNA insertion line (SALK_072730) in *FPA* (AT2G43410); homozygous plants were identified by three-primer PCR (Extended Data Fig. 2e,f).

For *pLD:LD-GFP* constructs, the full-length *LD* genomic sequence with its native promoter was fused to GFP in the *pCAMBIA1302* vector using the ClonExpress Entry One Step Cloning Kit (Vazyme, Cat. No. C112-01). The truncated *LD* genomic sequences were obtained by overlapping-PCR reactions and inserted into the *pCAMBIA1302* vector to generate *pLD:LD-ΔHOX-GFP*, which lacks the HOX domain (amino acids 63–129 deleted); *pLD:LD-ΔTFIIS-GFP*, which lacks the TFIIS domain (amino acids 231–324 deleted); and *pLD:LD-ΔC-GFP*, which lacks the C-terminal region (amino acids 501–953 deleted). For *pLD:LD-GFP-TurboID* proximity labelling constructs, the TurboID sequence was fused to the C-terminus of LD-GFP fusion proteins in a modified *pEarleyGate303* vector (the Gateway cassette replaced by a multiple cloning site) using the ClonExpress Entry One Step Cloning Kit. Truncation variants, including *pLD:LD-ΔTFIIS-GFP-TurboID* (lacking the TFIIS domain) and *pLD:LD-ΔHOX-GFP-TurboID* (lacking the HOX domain), were generated similarly. The above constructs were transformed into the *ld* mutant background. For the non-specific labelling control, GFP-TurboID was expressed under the control of the CaMV 35S promoter (*p35S:GFP-TurboID*) and transformed into the Col wild-type background.

For the *pFPA:FPA-GFP* construct, the full-length *FPA* genomic sequence with its native promoter was fused to GFP in the *pCAMBIA1302* vector and transformed into the *fpa-12* mutant background.

All constructs were introduced into *Agrobacterium tumefaciens* strain *GV3101* and transformed into the corresponding mutant backgrounds via the floral dip method^102^. Homozygous transgenic lines were obtained through antibiotic selection and genotyping after two generations.

### Estradiol-inducible aMIRFLD lines

To generate the *XVE::aMIRFLD* construct, the *pRS300* vector was used as a backbone to generate an artificial microRNA targeting *FLD*. Primers were designed according to WMD3 (http://wmd3.weigelworld.org/cgi-bin/webapp.cgi) and are listed in Supplementary Table 4. The aMIRFLD fragment was subcloned into the *pDONR221* vector, confirmed by sequencing, and recombined into the *pMDC7* destination vector wherein *aMIRFLD* expression is driven by an estradiol-inducible promoter. The construct was transformed into the Col wild-type background via floral dip. For induction experiments, seedlings or dry seeds were treated with 10 μM β-estradiol or DMSO control for the indicated durations.

### Chemical treatments

#### Flavopiridol treatment

For acute inhibition of RNAPII Ser2 phosphorylation, 12-day-old seedlings grown on solid medium were transferred to liquid 1/2 MS medium containing 5 μM flavopiridol (Cayman Chemical, Cat. No. 10009197) or an equivalent volume of DMSO. Seedlings were incubated with gentle agitation on an orbital shaker in conical flasks. Samples were collected at 15 min, 30 min, 1 h, and 2 h post-treatment for subsequent analyses.

#### 6-Azauracil (6-AU) treatment

To perturb transcription elongation dynamics, 12-day-old seedlings were transferred to liquid 1/2 MS medium containing 1.5 mg/L 6-azauracil (Sigma-Aldrich, A1757) or an equivalent volume of solvent control. Samples were collected at 12 h, 24 h, and 72 h post-treatment for time-course analyses of RNAPII redistribution, FLD relocalization, and H3K4me1 dynamics.

#### MG132 treatment

For proteasome inhibition experiments, 12-day-old seedlings grown on 1/2 MS medium were transferred to a new 9 × 9 cm Petri dish containing 50 mL of 50 μM MG132 (Abcam, Cat. No. ab141003), and then horizontally shaken at 60 rpm and 22°C for 24 h prior to sample collection.

### RNA isolation and RT-qPCR

Total RNA was extracted using the RNAprep Pure Plant Kit (MEGA, Cat. No. R4014-02) following the manufacturer’s instructions. First-strand cDNA was synthesized from 1 μg total RNA using HiScript II Q RT SuperMix (Vazyme, Cat. No. R223-01). Quantitative PCR was performed using SYBR Green Supermix (Vazyme, Cat. No. Q711-02) on a StepOnePlus Real-Time PCR System (Applied Biosystems). *UBQ10* was used as the internal reference gene. Relative expression levels were calculated using the 2^(-ΔΔCt) method^103^. Primers are listed in Supplementary Table 4. For *FLC* expression analysis, primers were designed to detect spliced (exon-spanning) and unspliced (intron-containing) transcripts separately.

### Protein extraction and immunoblotting

For nuclear protein extraction, approximately 2 g of 12-day-old seedlings were ground in liquid nitrogen and nuclei were isolated as described^104^. Nuclear proteins were extracted in lysis buffer (50 mM Tris-HCl [pH 8.0], 10 mM EDTA, 1% SDS, and protease inhibitors) at 4°C for 3 h. For total protein extraction, 0.2 g of 12-day-old seedlings were lysed in extraction buffer (50 mM Tris-HCl [pH 8.0], 300 mM NaCl, 10 mM EDTA, 0.1% Triton X-100, 10% glycerol, 0.2% NP-40, and protease inhibitors). Proteins were separated on 4–20% gradient SDS-PAGE gels (GenScript, SurePAGE) and transferred to PVDF membranes. Membranes were blocked in 5% non-fat milk and incubated with primary antibodies: anti-GFP (Abcam, Cat. No. ab290, 1:10,000), anti-RNAPII Ser2P (Abcam, Cat. No. ab193468, 1:10,000), anti-H3 (Proteintech, Cat. No. 17168-1-AP, 1:10,000). HRP-conjugated secondary antibodies and ECL detection were used for visualization.

### TurboID proximity labelling and mass spectrometry (TurboID-MS)

TurboID-based proximity labelling coupled with mass spectrometry was performed to identify protein-protein interactions^105,106^. Seedlings (12-day-old) expressing LD-GFP-TurboID variants, together with *p35S:GFP-TurboID* seedlings used as a non-specific labelling control, were treated with 50 μM biotin for 6 h before harvest. Approximately 0.7–1.0 g of biotin-treated plant material was ground in liquid nitrogen and lysed in 2.5 mL ice-cold lysis buffer (50 mM HEPES [pH 7.5], 300 mM NaCl, 10 mM EDTA, 10% glycerol, 0.2% NP-40, 1% Triton X-100, and 1× protease inhibitor cocktail) at 4°C with end-over-end rotation for 30–40 min. After sequential centrifugation (900 × *g* for 2 min, then 16,000 × *g* for 10 min, twice), the clarified lysate was subjected to buffer exchange using PD-10 desalting columns (Cytiva, Cat. No. 17085101). Biotinylated proteins were captured using streptavidin-conjugated magnetic beads (MCE, Cat. No. HY-K0208, 50 μL) at 4°C for 4 h with end-over-end rotation. Beads were washed four times with wash buffer (50 mM HEPES [pH 7.5], 100 mM NaCl, 10 mM EDTA, 10% glycerol, 0.1% NP-40, and 1× protease inhibitor cocktail). On-bead digestion was performed by resuspending beads in 100 μL elution buffer I (50 mM Tris-HCl [pH 7.5], 2 M urea, 1 mM DTT) with 2 μL sequencing-grade trypsin at 30°C for 1.5 h, followed by addition of 400 μL elution buffer II (50 mM Tris-HCl [pH 7.5], 2 M urea, 5 mM iodoacetamide) and overnight digestion at 32°C. Peptides were desalted using C18 spin columns and peptides were extracted with 5% formic acid/50% acetonitrile, dried, and redissolved in 0.1% formic acid. Samples were analysed on a Q Exactive HF mass spectrometer (Thermo Fisher) coupled with an Ultimate 3000 UHPLC system in data-dependent acquisition (DDA) mode, where the top 20 most abundant precursor ions were selected for fragmentation analysis. Mobile phase A consisted of 0.1% formic acid solution, while mobile phase B was 80% acetonitrile containing 0.1% formic acid. The flow rate was set at 400 nL/min for the Nano pump and 10 μL/min for the loading pump.

The Nano Trap column (Lot No. 164535, Bed Length = 20 mm) and the analytical column (Lot No. 164941, Length = 250 mm) were provided by Thermo Fisher Scientific. RAW files were processed with MaxQuant (v2.4.2.0) using the built-in Andromeda search engine, searching against the Arabidopsis thaliana TAIR10 protein database supplemented with the GFP and TurboID sequences.

Differential enrichment analysis compared each LD truncation variant (LD-ΔHOX, LD-ΔTFIIS) with full-length LD-GFP-TurboID as the matched reference, ensuring that all samples shared the same TurboID fusion architecture and expression driver. Proteins showing more than fourfold enrichment with P < 0.05 were considered significantly enriched.

### Recombinant protein expression and purification

Recombinant FLD and LD fragments used for in vitro interaction assays were cloned into bacterial expression vectors to generate GST-or MBP–His-tagged fusion proteins. These comprised a fragment within the FLD C-terminal region (FLD-C; amino acids 672–842) and the LD HOX- and TFIIS-containing fragments. Proteins were expressed in *Escherichia coli* BL21 (DE3) and induced with 0.5 mM IPTG at 18°C for 16 h. GST-tagged proteins were purified using glutathione-Sepharose beads (Beyotime, Cat. No. P2262), and MBP–His-tagged proteins were purified using amylose resin (Beyotime, Cat. No. P2210) according to the manufacturer’s instructions. Protein purity was assessed by SDS–PAGE and Coomassie staining.

### CTD peptide pull-down assays

Synthetic biotinylated CTD peptides containing either unphosphorylated YSPTSPS or Ser2-phosphorylated YpSPTSPS heptad repeats were custom synthesized by GenScript. Peptides (1 μg each) were immobilized on Dynabeads MyOne Streptavidin T1 (Thermo Fisher, Cat. No. 65601) by incubation for 1 h at room temperature. Beads were washed and incubated with purified MBP-HIS-LD-TFIIS proteins (1 μg) in binding buffer (30 mM HEPES [pH 7.5], 150 mM NaCl, 0.1% NP-40) at 4°C for 2 h. After extensive washing, bound proteins were eluted, separated by SDS-PAGE, and detected by immunoblotting with anti-MBP antibody (TransGen Biotech, Cat. No. HT701-01, 1:3,000).

### *In vitro* pull-down assays

For testing direct protein-protein interactions between the FLD C terminus and LD domains, GST-tagged FLD-C was incubated with MBP-HIS-tagged LD-HOX or LD-TFIIS domains in binding buffer (30 mM HEPES [pH 7.5], 150 mM NaCl, 0.1% NP-40) at 4°C for 2 h. Complexes were captured on glutathione beads, washed extensively, and analysed by immunoblotting. The proteins were immunoblotted with anti-GST (TransGen Biotech, Cat. No. HT601-01, 1:5,000) or anti-MBP (TransGen Biotech, Cat. No. HT701-01, 1:3,000) primary antibodies, followed by horseradish peroxidase-conjugated goat anti-mouse secondary antibody (Abcam, Cat. No. ab6789, 1:10,000).

### Structural prediction of FLD and FLD-containing complexes

For prediction of the FLD-containing complexes, AlphaFold 3 was used with default settings^107,108^. Because the FLD complex contains extended intrinsically disordered regions, interface quality was evaluated using interface-residue-pair PAE values rather than whole-complex ipTM scores, which are biased by disorder-related uncertainty. Predicted structures were visualized in PyMOL, and interface residues and predicted contacts were analyzed on the basis of the top-ranked models.

### ChIP-qPCR and ChIP-seq

ChIP experiments were performed as previously described with minor modifications^109,110^. Briefly, 1 g of 12-day-old seedlings (per biological replicate) were crosslinked with 1% formaldehyde under vacuum for 15 min and quenched with 0.125 M glycine. Crosslinked tissue was ground to fine powder in liquid nitrogen. Chromatin was extracted and sonicated to 200–500 bp fragments using a Bioruptor sonicator (Diagenode) with 30 s on/30 s off cycles (27 total on cycles) at the HIGH setting. Sonicated chromatin was diluted 10-fold with ChIP dilution buffer (16.7 mM Tris-HCl [pH 8.0], 167 mM NaCl, 1.1% Triton X-100, 1.2 mM EDTA) and incubated overnight at 4°C with the following antibodies: anti-GFP (Abcam, Cat. No. ab290, 1 μL), anti-RNAPII Ser2P (Abcam, Cat. No. ab193468, 1 μL), anti-H3K4me1 (Abcam, Cat. No. ab8895, 1 μL). Immune complexes were captured with Magna ChIP Protein A magnetic beads (Merck, Cat. No. 16-661), washed sequentially with low salt, high salt, LiCl, and TE buffers, and eluted in elution buffer (1% SDS, 0.1 M NaHCO₃). After reverse crosslinking at 65°C overnight and treatment with proteinase K, DNA was purified using the MinElute PCR Purification Kit (Qiagen, Cat. No. 28004).

For ChIP-qPCR, results were calculated as a percentage of input DNA according to the Champion ChIP-qPCR user manual (SABiosciences). Primers used for ChIP-qPCR are listed in Supplementary Table 4. ChIP-seq libraries were prepared from 1–2 ng of ChIP DNA using the VAHTS Universal DNA Library Prep Kit for Illumina V3 (Vazyme, Cat. No. ND607), VAHTS DNA Adapters set3–set6 for Illumina (Vazyme, Cat. No. N805), and VAHTS DNA Clean Beads (Vazyme, Cat. No. N411-02) following the manufacturer’s instructions. Libraries were sequenced on the Illumina NovaSeq platform with paired-end 150 bp reads. Two biological replicates were performed for each condition.

### ChIP-seq data analysis

ChIP−seq data were analysed following previously established methods^111^. In brief, raw reads were aligned to the *Arabidopsis thaliana* reference genome (TAIR10) using Bowtie2 with default settings^112^. Only uniquely mapped reads were retained for subsequent analysis. MACS2 was employed to call peaks using the following parameters: “gsize = 119,667,750, bw = 300, q = 0.05, nomodel, extsize = 200.” Peaks present in both biological replicates (irreproducible discovery rate < 0.05) were retained for further analysis^113^. ChIPseeker was utilised to annotate peaks to genes with default settings^114^. Differential binding analysis was performed using DiffBind with default settings^115^; full results for all pairwise comparisons are provided in Supplementary Table 2. Venn diagrams were created using the Venny website (https://bioinfogp.cnb.csic.es/tools/venny/index2.0.2.html). The total number of genes in the *Arabidopsis* genome (TAIR10) is 34,218. For visualization and downstream analysis, bigwig files were generated by bamCoverage with “bin size 10” and “normalize to RPKM (reads per kilobase per million)” in deepTools and visually displayed by Integrative Genomics Viewer (IGV)^116^. Metagene profiles were generated using computeMatrix and plotProfile functions in deepTools. To analyse read density and correlation between different ChIP−seq samples, Spearman correlation analysis was performed. Read density was analysed over the merged set of binding sites across all ChIPs using the multiBigwigSummary function in bin mode (bin size = 1,000), and the heatmap of Spearman correlation was generated by the plotCorrelation function in deepTools^117^.

### Plant Native Elongating Transcript sequencing (pNET-seq)

pNET-seq was performed in 12-day-old seedlings as previously described with modifications^118,119^. For the 6-AU time-course experiments, seedlings were treated with 6-AU as described above, and samples were collected at the indicated time points. Briefly, approximately 2 g of seedlings were crosslinked with 37 mL of 1% formaldehyde under vacuum in two rounds (10 min and 20 min). The crosslinking was quenched by adding 2.5 mL of 2 M glycine for 5 min. Crosslinked tissue was ground into a fine powder in liquid nitrogen. The tissue powder was solubilized in 30 mL of nuclei lysis buffer (50 mM HEPES [pH 7.5], 150 mM NaCl, 1 mM EDTA, 1% (w/v) Triton X-100, 10% (w/v) glycerol, 5 mM 2-mercaptoethanol, 1× protease inhibitor cocktail, 1 mM PMSF) and filtered through two layers of Miracloth. The nuclei were spun down at 3,200 × g for 20 min at 4°C, then washed once separately with HBB buffer (25 mM Tris-HCl [pH 7.5], 0.44 M sucrose, 10 mM MgCl₂, 0.1% (w/v) Triton X-100, 10 mM 2-mercaptoethanol) and HBC buffer (20 mM Tris-HCl [pH 7.5], 352 mM sucrose, 8 mM MgCl₂, 0.08% (w/v) Triton X-100, 8 mM 2-mercaptoethanol, 20% (w/v) glycerol).

The nuclei pellets were resuspended in 400 μL MNase buffer (20 mM Tris-HCl [pH 8.0], 5 mM NaCl, 2.5 mM CaCl₂) and digested with 20 U MNase (TaKaRa) for 5 min at 37°C with rotation at 1,400 rpm. The digestion was stopped by adding 16 μL of 500 mM EDTA. The digested chromatin was subjected to mild sonication to release the RNAPII and nascent RNA complex. After centrifugation at 16,000 × g for 5 min at 4°C, the supernatant was collected and diluted 10-fold with IP buffer, followed by overnight immunoprecipitation with anti-Ser2P CTD RNAPII antibody (Abcam, Cat. No. ab5095) and Protein A Dynabeads (Thermo Fisher Scientific, Cat. No. 10002D). After immunoprecipitation, the beads were washed 6 times with NET-2 buffer (10 mM Tris-HCl [pH 7.5], 150 mM NaCl, 0.05% (v/v) NP-40), followed by one wash with 1× PNK buffer containing 0.1% Triton X-100. The resulting nascent RNA bound to RNAPII was treated with 50 μL PNK reaction mix (5 μL 10× PNK buffer, 5 μL 10 mM ATP, 1 μL T4 PNK (Thermo Fisher Scientific), 39 μL H₂O) on the beads at 37°C for 8 min for RNA 5’ end phosphorylation. After washing once with NET-2 buffer, the beads were eluted twice with 55 μL Elution Buffer containing RNase inhibitor (RiboLock) at room temperature for 10 min and 65°C for 15 min, respectively. The combined eluate was treated with 20 μg RNA-grade Proteinase K at 55°C for 1 h. The resulting RNA was extracted using TRIzol and precipitated with isopropanol in the presence of glycogen overnight.

The sequencing library was constructed using the VAHTS Small RNA Library Prep Kit for Illumina (Vazyme, Cat. No. NR811) according to the manufacturer’s instructions. Libraries were sequenced on the Illumina NovaSeq platform with paired-end 150 bp reads. Three biological replicates were performed for each condition.

### pNET-seq data analysis

pNET-seq data were analysed following previously established methods with modifications^118^. Briefly, adapter sequences were removed using cutadapt (v3.4), reads were subsampled with seqtk to a common depth determined by the smallest retained library size, and aligned to the *Arabidopsis thaliana* reference genome (TAIR10) using STAR. After filtering for properly paired reads and removing reads mapping to rRNA, snRNA, tRNA and snoRNA, strand-specific coverage tracks were generated from the 3′ end position of each read to capture the RNAPII active site. *FLC*-specific transcriptional metrics, including 5′ signal ratio, termination index and elongation-speed estimation, were calculated as described below. Custom code used for transcription-speed estimation from pNET-seq data is described in the Code availability statement.

### Quantification of *FLC* transcriptional metrics from pNET-seq

For *FLC* locus analysis, strand-specific pNET-seq signal was quantified over defined intervals relative to annotated transcription landmarks. The sense-strand 5′ signal ratio was calculated as the fraction of signal within a 1.5-kb TSS-proximal window relative to total signal across the annotated *FLC* transcript. The antisense 5′ signal ratio was calculated analogously using a 500-bp window downstream of the *COOLAIR* TSS, because *COOLAIR* comprises multiple isoforms with heterogeneous 3′ ends, precluding a uniform 1.5-kb window definition.

The termination index was adapted from a previously described metric^88^ and was calculated here as the ratio of sense-strand pNET-seq signal in a termination window spanning 150 bp upstream to 400 bp downstream of the annotated transcription termination site (TTS) to signal in the remaining gene body. For *FLC*, upstream and downstream were defined relative to the direction of transcription.

RNAPII elongation speed across *FLC* intron 1 was estimated from sense-strand pNET-seq signal using the reverse cumulative sum approach. Briefly, pNET-seq signal across the analysed intronic region was divided into consecutive 10-bp bins, and a reverse cumulative sum was calculated from the 3′ end of the intron towards the 5′ end in the direction of transcription. The slope of a linear regression fitted to the cumulative signal profile was used to derive an elongation-speed metric, expressed as log2(−1/slope). The implementation used for these calculations is provided in the custom analysis code.

### Flowering time measurements

Flowering time was measured as the total number of rosette leaves at bolting (when the primary inflorescence reached 1 cm in height). Plants were grown under long-day conditions (16 h light/8 h dark) at 22°C. At least 10 individual plants were scored for each genotype.

### FLD homologues alignments

Full-length amino acid sequences of FLD and representative LSD1-family homologues across plants, metazoans and fungi were retrieved from NCBI. Multiple sequence alignment was performed with ClustalW as implemented in MEGA7 using default parameters. A phylogenetic tree was constructed by the neighbour-joining method with 1,000 bootstrap replicates. The resulting tree was visualized and annotated using iTOL.

### Statistics and reproducibility

Unless otherwise stated in the relevant figure legend, data in bar graphs and line graphs are presented as mean ± s.d. For violin plots, solid lines indicate the median, dashed lines indicate the upper and lower quartiles, and outliers were determined using the Tukey method. For box plots, the centre line indicates the median, box limits indicate the interquartile range (IQR), and whiskers extend to 1.5 × IQR. Unless otherwise stated, statistical significance for experiments with three or more groups was assessed by one-way ANOVA with Tukey’s post hoc test, and different letters denote statistically significant differences at *P* < 0.05. For two-group comparisons, a two-tailed Student’s *t*-test was used. Asterisks denote \**P* < 0.05, \*\**P* < 0.01, \*\*\**P* < 0.001 and \*\*\*\**P* < 0.0001; ns, not significant. The number of biological replicates (n) is indicated in the corresponding figure legends. No statistical methods were used to predetermine sample sizes. No data were excluded from analyses. The experiments were not randomised. The investigators were not blinded to allocation during experiments and outcome assessment.

### Software used in this study

Data analyses were performed with the following software:

Sequencing data processing: cutadapt (v3.4), seqtk (v1.3), Bowtie2 (v2.5.0), STAR (v2.7.10a), samtools (v1.17).

ChIP-seq analysis: MACS2 (v2.2.7.1), ChIPseeker (v1.36.0), DiffBind (v3.10.0), deepTools (v3.5.1).

Mass spectrometry: MaxQuant (v2.4.2.0) with the Andromeda search engine; Perseus (v2.0.5.0).

Visualization and structural analysis: IGV (v2.16.0), PyMOL (v2.5.5), Fiji/ImageJ (v2.14.0), GraphPad Prism (v9.5.1).

Structural prediction: AlphaFold 3 (default parameters).

## ACKNOWLEDGMENT

We thank the Arabidopsis Biological Resource Center (ABRC) for T-DNA insertion lines, Prof. Hongchun Yang (Wuhan University) for providing the *FRI-Col* line, and Prof. Xin-Jian He (National Institute of Biological Sciences, Beijing) for providing *ld* mutant. We thank the Mass Spectrometry Platform at Sun Yat-sen University for LC-MS/MS analysis. This work was supported by the National Natural Science Foundation of China to C.L. (32470346, 32670388), and the National Key Research and Development Program of China (2024YFD1200800).

## AUTHOR CONTRIBUTIONS

C.L. and X.S. conceived the project and designed the research. X.S. performed most of the experiments. X.S., Y.L., C.C. and C.L. conducted bioinformatics analysis. X.S. and L.Y. generated the plant materials and constructs. X.S., Y.L., L.Y., Y.H., J.L., Y.Z., H.H., Y.H., Z.Z., Z.L., T.Z., C.C. and C.L. analyzed data. C.L. and X.S. wrote the manuscript.

## Data availability

ChIP-seq data generated in this study have been deposited in the Gene Expression Omnibus (GEO) database under accession number GSE327776. pNET-seq data have been deposited under accession number GSE341142. TurboID-MS proteomics data have been deposited in the iProX database under the dataset identifier IPX0016621000. Any additional information required to reanalyze the data reported in this paper is available from the lead contact upon request.

## Code availability

Custom code used for transcription-speed estimation from pNET-seq data is available from the corresponding author upon reasonable request. All other analyses were performed using published software as described in the Methods.

**Supplementary Table 1.** RNAPII-Ser2P and FLD ChIP-seq target gene sets and their convergently oriented subsets.

**Supplementary Table 2.** Differential binding analyses of FLD and LD ChIP-seq data.

**Supplementary Table 3.** Proteins identified by TurboID-based proximity labelling of full-length LD, relative to the p35S:GFP-TurboID.

**Supplementary Table 4.** Oligonucleotides used in this study.

## References

1 Li, B., Carey, M. & Workman, J. L. The role of chromatin during transcription. Cell 128, 707–719, doi:10.1016/j.cell.2007.01.015 (2007).

2 Peterson, C. L. & Laniel, M.-A. Histones and histone modifications. Curr. Biol. 14, R546–R551, doi:10.1016/j.cub.2004.07.007 (2004).

3 Strahl, B. D. & Allis, C. D. The language of covalent histone modifications. Nature 403, 41–45 (2000).

4 Klemm, S. L., Shipony, Z. & Greenleaf, W. J. Chromatin accessibility and the regulatory epigenome. Nat. Rev. Genet. 20, 207–220, doi:10.1038/s41576-018-0089-8 (2019).

5 Thurman, R. E. et al. The accessible chromatin landscape of the human genome. Nature 489, 75–82, doi:10.1038/nature11232 (2012).

6 Guertin, M. J. & Lis, J. T. Mechanisms by which transcription factors gain access to target sequence elements in chromatin. Curr. Opin. Genet. Dev. 23, 116–123, doi:10.1016/j.gde.2012.11.008 (2013).

7 Eberharter, A. & Becker, P. B. Histone acetylation: a switch between repressive and permissive chromatin. EMBO Rep. 3, 224–229 (2002).

8 Clayton, A. L., Hazzalin, C. A. & Mahadevan, L. C. Enhanced histone acetylation and transcription: a dynamic perspective. Mol. Cell 23, 289–296, doi:10.1016/j.molcel.2006.06.017 (2006).

9 Kouzarides, T. Chromatin modifications and their function. Cell 128, 693–705, doi:10.1016/j.cell.2007.02.005 (2007).

10 Agalioti, T., Chen, G. & Thanos, D. Deciphering the transcriptional histone acetylation code for a human gene. Cell 111, 381–392 (2002).

11 Barski, A. et al. High-resolution profiling of histone methylations in the human genome. Cell 129, 823–837, doi:10.1016/j.cell.2007.05.009 (2007).

12 Benayoun, B. A. et al. H3K4me3 breadth is linked to cell identity and transcriptional consistency. Cell 158, 673–688, doi:10.1016/j.cell.2014.06.027 (2014).

13 Soares, L. M. et al. Determinants of histone H3K4 methylation patterns. Mol. Cell 68, 773–785, doi:10.1016/j.molcel.2017.10.013 (2017).

14 Wang, H. et al. H3K4me3 regulates RNA polymerase II promoter-proximal pause-release. Nature 615, 339–348, doi:10.1038/s41586-023-05780-8 (2023).

15 Shilatifard, A. The COMPASS family of histone H3K4 methylases: mechanisms of regulation in development and disease pathogenesis. Annu. Rev. Biochem. 81, 65–95, doi:10.1146/annurev-biochem-051710-134100 (2012).

16 Santos-Rosa, H. et al. Active genes are tri-methylated at K4 of histone H3. Nature 419, 403–407, doi:10.1038/nature01071 (2002).

17 Kim, T. & Buratowski, S. Dimethylation of H3K4 by Set1 recruits the Set3 histone deacetylase complex to 5′ transcribed regions. Cell 137, 259–272, doi:10.1016/j.cell.2009.02.045 (2009).

18 Krogan, N. J. et al. Methylation of histone H3 by Set2 in *Saccharomyces cerevisiae* is linked to transcriptional elongation by RNA polymerase II. Mol. Cell. Biol. 23, 4207–4218, doi:10.1128/MCB.23.12.4207-4218.2003 (2003).

19 Schaft, D. et al. The histone 3 lysine 36 methyltransferase, SET2, is involved in transcriptional elongation. Nucleic Acids Res. 31, 2475-2482, doi:10.1093/nar/gkg372 (2003).

20 Kizer, K. O. et al. A novel domain in Set2 mediates RNA polymerase II interaction and couples histone H3 K36 methylation with transcript elongation. Mol. Cell. Biol. 25, 3305–3316, doi:10.1128/MCB.25.8.3305-3316.2005 (2005).

21 Wagner, E. J. & Carpenter, P. B. Understanding the language of Lys36 methylation at histone H3. Nat. Rev. Mol. Cell Biol. 13, 115–126, doi:10.1038/nrm3274 (2012).

22 Gopalakrishnan, R., Marr, S. K., Kingston, R. E. & Winston, F. A conserved genetic interaction between Spt6 and Set2 regulates H3K36 methylation. Nucleic Acids Res. 47, 3888–3903, doi:10.1093/nar/gkz119 (2019).

23 Keogh, M. C. et al. Cotranscriptional set2 methylation of histone H3 lysine 36 recruits a repressive Rpd3 complex. Cell 123, 593–605, doi:10.1016/j.cell.2005.10.025 (2005).

24 Carrozza, M. J. et al. Histone H3 methylation by Set2 directs deacetylation of coding regions by Rpd3S to suppress spurious intragenic transcription. Cell 123, 581–592, doi:10.1016/j.cell.2005.10.023 (2005).

25 Riedl, T. & Egly, J.-M. Phosphorylation in transcription: the CTD and more. Gene Expr. 9, 3–13 (2000).

26 Ho, C. K. & Shuman, S. Distinct roles for CTD Ser-2 and Ser-5 phosphorylation in the recruitment and allosteric activation of mammalian mRNA capping enzyme. Mol. Cell 3, 405–411 (1999).

27 Buratowski, S. Progression through the RNA polymerase II CTD cycle. Mol. Cell 36, 541–546, doi:10.1016/j.molcel.2009.10.019 (2009).

28 Hajheidari, M., Koncz, C. & Eick, D. Emerging roles for RNA polymerase II CTD in *Arabidopsis*. Trends Plant Sci. 18, 633–643, doi:10.1016/j.tplants.2013.07.001 (2013).

29 Cramer, P. Eukaryotic transcription turns 50. Cell 179, 808–812, doi:10.1016/j.cell.2019.09.018(2019).

30 Hsin, J.-P. & Manley, J. L. The RNA polymerase II CTD coordinates transcription and RNA processing. Genes Dev. 26, 2119–2137, doi:10.1101/gad.200303.112 (2012).

31 Margueron, R. & Reinberg, D. The Polycomb complex PRC2 and its mark in life. Nature 469, 343–349 (2011).

32 Schuettengruber, B., Bourbon, H. M., Di Croce, L. & Cavalli, G. Genome regulation by Polycomb and Trithorax: 70 years and counting. Cell 171, 34–57, doi:10.1016/j.cell.2017.08.002 (2017).

33 D’Urso, A. & Brickner, J. H. Mechanisms of epigenetic memory. Trends Genet. 30, 230–236 (2014).

34 Vermunt, M. W. et al. Gene silencing dynamics are modulated by transiently active regulatory elements. Mol. Cell 83, 715–730, doi:10.1016/j.molcel.2023.02.006 (2023).

35 Wutz, A. Gene silencing in X-chromosome inactivation: advances in understanding facultative heterochromatin formation. Nat. Rev. Genet. 12, 542–553, doi:10.1038/nrg3035 (2011).

36 Reik, W. Stability and flexibility of epigenetic gene regulation in mammalian development. Nature 447, 425–432, doi:10.1038/nature05918 (2007).

37 Montez, M., Schulten, A., Menon, G. & Dean, C. Molecular integration of seasonal temperature signals in flowering time control. Nat. Rev. Genet. 27, 695–707, doi:10.1038/s41576-026-00979-7 (2026).

38 Verdel, A. et al. RNAi-mediated targeting of heterochromatin by the RITS complex. Science 303, 672–676, doi:10.1126/science.1093686 (2004).

39 Moazed, D. Small RNAs in transcriptional gene silencing and genome defence. Nature 457, 413–420 (2009).

40 Aravin, A. A., Sachidanandam, R., Girard, A., Fejes-Toth, K. & Hannon, G. J. Developmentally regulated piRNA clusters implicate MILI in transposon control. Science 316, 744–747, doi:10.1126/science.1142612 (2007).

41 Houwing, S. et al. A role for Piwi and piRNAs in germ cell maintenance and transposon silencing in zebrafish. Cell 129, 69–82, doi:10.1016/j.cell.2007.03.026 (2007).

42 Khurana, J. S. & Theurkauf, W. piRNAs, transposon silencing, and *Drosophila* germline development. J. Cell Biol. 191, 905–913, doi:10.1083/jcb.201006034 (2010).

43 Sienski, G., Donertas, D. & Brennecke, J. Transcriptional silencing of transposons by Piwi and maelstrom and its impact on chromatin state and gene expression. Cell 151, 964–980, doi:10.1016/j.cell.2012.10.040 (2012).

44 Zhao, J., Sun, B. K., Erwin, J. A., Song, J.-J. & Lee, J. T. Polycomb proteins targeted by a short repeat RNA to the mouse X-chromosome. Science 322, 750–756, doi:10.1126/science.1163045 (2008).

45 Pintacuda, G. et al. hnRNPK recruits PCGF3/5-PRC1 to the Xist RNA B-repeat to establish polycomb-mediated chromosomal silencing. Mol. Cell 68, 955–969, doi:10.1016/j.molcel.2017.11.013 (2017).

46 Churchman, L. S. & Weissman, J. S. Nascent transcript sequencing visualizes transcription at nucleotide resolution. Nature 469, 368–373, doi:10.1038/nature09652 (2011).

47 Adelman, K. & Lis, J. T. Promoter-proximal pausing of RNA polymerase II: emerging roles in metazoans. Nat. Rev. Genet. 13, 720–731, doi:10.1038/nrg3293 (2012).

48 Henriques, T. et al. Stable pausing by RNA polymerase II provides an opportunity to target and integrate regulatory signals. Mol. Cell 52, 517–528, doi:10.1016/j.molcel.2013.10.001 (2013).

49 Imashimizu, M. et al. Intrinsic translocation barrier as an initial step in pausing by RNA polymerase II. J. Mol. Biol. 425, 697–712, doi:10.1016/j.jmb.2012.12.002 (2013).

50 Kwak, H., Fuda, N. J., Core, L. J. & Lis, J. T. Precise maps of RNA polymerase reveal how promoters direct initiation and pausing. Science 339, 950–953, doi:10.1126/science.1229386 (2013).

51 Weber, C. M., Ramachandran, S. & Henikoff, S. Nucleosomes are context-specific, H2A.Z-modulated barriers to RNA polymerase. Mol. Cell 53, 819–830, doi:10.1016/j.molcel.2014.02.014 (2014).

52 Kwak, H. & Lis, J. T. Control of transcriptional elongation. Annu. Rev. Genet. 47, 483–508, doi:10.1146/annurev-genet-110711-155440 (2013).

53 Jonkers, I. & Lis, J. T. Getting up to speed with transcription elongation by RNA polymerase II. Nat. Rev. Mol. Cell Biol. 16, 167–177, doi:10.1038/nrm3953 (2015).

54 Noe Gonzalez, M., Blears, D. & Svejstrup, J. Q. Causes and consequences of RNA polymerase II stalling during transcript elongation. Nat. Rev. Mol. Cell Biol. 22, 3–21, doi:10.1038/s41580-020-00308-8 (2021).

55 Gaertner, B. & Zeitlinger, J. RNA polymerase II pausing during development. Development 141, 1179–1183, doi:10.1242/dev.088492 (2014).

56 Core, L. & Adelman, K. Promoter-proximal pausing of RNA polymerase II: a nexus of gene regulation. Genes Dev. 33, 960–982, doi:10.1101/gad.325142.119 (2019).

57 Levine, M. Paused RNA polymerase II as a developmental checkpoint. Cell 145, 502–511, doi:10.1016/j.cell.2011.04.021 (2011).

58 Heintzman, N. D. et al. Distinct and predictive chromatin signatures of transcriptional promoters and enhancers in the human genome. Nature Genet. 39, 311–318, doi:10.1038/ng1966 (2007).

59 Heintzman, N. D. et al. Histone modifications at human enhancers reflect global cell-type-specific gene expression. Nature 459, 108–112, doi:10.1038/nature07829 (2009).

60 Zhang, X., Bernatavichute, Y. V., Cokus, S., Pellegrini, M. & Jacobsen, S. E. Genome-wide analysis of mono-, di- and trimethylation of histone H3 lysine 4 in *Arabidopsis thaliana*. Genome Biol. 10, R62, doi:10.1186/gb-2009-10-6-r62 (2009).

61 Pokholok, D. K. et al. Genome-wide map of nucleosome acetylation and methylation in yeast. Cell 122, 517–527, doi:10.1016/j.cell.2005.06.026 (2005).

62 Oya, S., Takahashi, M., Takashima, K., Kakutani, T. & Inagaki, S. Transcription-coupled and epigenome-encoded mechanisms direct H3K4 methylation. Nature communications 13, 4521, doi:10.1038/s41467-022-32165-8 (2022).

63 Wu, J. et al. H3K4me1 directs H3K36me2 and H3K36me3 deposition in land plants. Nature communications 17, 2831, doi:10.1038/s41467-026-69632-5 (2026).

64 Tao, L. et al. Enhancer decommissioning imposes an epigenetic barrier to sensory hair cell regeneration. Dev. Cell 56, 2471–2485, doi:10.1016/j.devcel.2021.07.003 (2021).

65 Inagaki, S. et al. Gene-body chromatin modification dynamics mediate epigenome differentiation in *Arabidopsis*. EMBO J. 36, 970–980, doi:10.15252/embj.201694983 (2017).

66 Vinckier, N. K. et al. LSD1-mediated enhancer silencing attenuates retinoic acid signalling during pancreatic endocrine cell development. Nat. Commun. 11, 2082, doi:10.1038/s41467-020-16017-x (2020).

67 Whyte, W. A. et al. Enhancer decommissioning by LSD1 during embryonic stem cell differentiation. Nature 482, 221–225, doi:10.1038/nature10805 (2012).

68 He, Y., Michaels, S. D. & Amasino, R. M. Regulation of flowering time by histone acetylation in *Arabidopsis*. Science 302, 1751–1754, doi:10.1126/science.1091109 (2003).

69 Inagaki, S., Takahashi, M., Takashima, K., Oya, S. & Kakutani, T. Chromatin-based mechanisms to coordinate convergent overlapping transcription. Nat. Plants 7, 295–302, doi:10.1038/s41477-021-00868-3 (2021).

70 Liu, F. et al. The *Arabidopsis* RNA-binding protein FCA requires a lysine-specific demethylase 1 homolog to downregulate *FLC*. Mol. Cell 28, 398–407, doi:10.1016/j.molcel.2007.10.018 (2007).

71 Fang, X. et al. The 3′ processing of antisense RNAs physically links to chromatin-based transcriptional control. Proc. Natl Acad. Sci. USA 117, 15316–15321, doi:10.1073/pnas.2007268117 (2020).

72 Mateo-Bonmatí, E. et al. A CPF-like phosphatase module links transcription termination to chromatin silencing. Mol. Cell 84, 2272–2286, doi:10.1016/j.molcel.2024.05.016 (2024).

73 Rahl, P. B. et al. c-Myc regulates transcriptional pause release. Cell 141, 432–445, doi:10.1016/j.cell.2010.03.030 (2010).

74 Mayer, A. et al. Uniform transitions of the general RNA polymerase II transcription complex. Nat. Struct. Mol. Biol. 17, 1272–1278, doi:10.1038/nsmb.1903 (2010).

75 Zhu, J., Liu, M., Liu, X. & Dong, Z. RNA polymerase II activity revealed by GRO-seq and pNET-seq in *Arabidopsis*. Nat. Plants 4, 1112–1123, doi:10.1038/s41477-018-0280-0 (2018).

76 Yu, X., Martin, P. G. P. & Michaels, S. D. BORDER proteins protect expression of neighboring genes by promoting 3′ Pol II pausing in plants. Nat. Commun. 10, 4359, doi:10.1038/s41467-019-12328-w (2019).

77 Mayer, A. et al. Native elongating transcript sequencing reveals human transcriptional activity at nucleotide resolution. Cell 161, 541–554, doi:10.1016/j.cell.2015.03.010 (2015).

78 Chen, C. et al. RNA polymerase II-independent recruitment of SPT6L at transcription start sites in Arabidopsis. Nucleic Acids Res. 47, 6714–6725, doi:10.1093/nar/gkz465 (2019).

79 Macknight, R., et al. *FCA*, a gene controlling flowering time in *Arabidopsis*, encodes a protein containing RNA-binding domains. Cell 89, 737–745 (1997).

80 Hornyik, C., Terzi, L. C. & Simpson, G. G. The spen family protein FPA controls alternative cleavage and polyadenylation of RNA. Dev. Cell 18, 203–213, doi:10.1016/j.devcel.2009.12.009 (2010).

81 Wu, Z., Fang, X., Zhu, D. & Dean, C. Autonomous pathway: *FLOWERING LOCUS C* repression through an antisense-mediated chromatin-silencing mechanism. Plant Physiol. 182, 27–37, doi:10.1104/pp.19.01009 (2020).

82 Schomburg, F. M., Patton, D. A., Meinke, D. W. & Amasino, R. M. FPA, a gene involved in floral induction in *Arabidopsis*, encodes a protein containing RNA-recognition motifs. Plant Cell 13, 1427–1436, doi:10.1105/tpc.13.6.1427 (2001).

83 Liu, F., Marquardt, S., Lister, C., Swiezewski, S. & Dean, C. Targeted 3′ processing of antisense transcripts triggers *Arabidopsis FLC* chromatin silencing. Science 327, 94–97 (2010).

84 Johanson, U. et al. Molecular analysis of *FRIGIDA*, a major determinant of natural variation in *Arabidopsis* flowering time. Science 290, 344–347 (2000).

85 Exinger, F. & Lacroute, F. 6-Azauracil inhibition of GTP biosynthesis in *Saccharomyces cerevisiae*. Curr. Genet. 22, 9–11 (1992).

86 Whittaker, C. & Dean, C. The *FLC* locus: a platform for discoveries in epigenetics and adaptation. Annu. Rev. Cell Dev. Biol. 33, 555–575, doi:10.1146/annurev-cellbio- (2017).

87 Yang, H. et al. Distinct phases of Polycomb silencing to hold epigenetic memory of cold in *Arabidopsis*. Science 357, 1142–1145 (2017).

88 Maple, R. et al. Co-transcriptional splicing changes combine with reduced productive transcription initiation for cold-induced repression of *FLC*. Nucleic Acids Res. 54, gkaf1462, doi:10.1093/nar/gkaf1462 (2026).

89 Qi, P.-L. et al. Characterization of an autonomous pathway complex that promotes flowering in *Arabidopsis*. Nucleic Acids Res. 50, 7380–7395, doi:10.1093/nar/gkac551 (2022).

90 Mori, S. et al. Cotranscriptional demethylation induces global loss of H3K4me2 from active genes in *Arabidopsis*. EMBO J. 42, e113798, doi:10.15252/embj.2023113798 (2023).

91 Baxter, C. L., Šviković, S., Sale, J. E., Dean, C. & Costa, S. The intersection of DNA replication with antisense 3′ RNA processing in *Arabidopsis FLC* chromatin silencing. Proc. Natl Acad. Sci. USA 118, e2107483118, doi:10.1073/pnas.2107483118 (2021).

92 Angel, A., Song, J., Dean, C. & Howard, M. A Polycomb-based switch underlying quantitative epigenetic memory. Nature 476, 105–108, doi:10.1038/nature10241 (2011).

93 Menon, G., Schulten, A., Dean, C. & Howard, M. Digital paradigm for Polycomb epigenetic switching and memory. Curr. Opin. Plant Biol. 61, 102012, doi:10.1016/j.pbi.2021.102012 (2021).

94 Antoniou-Kourounioti, R. L. et al. Integrating analog and digital modes of gene expression at *Arabidopsis FLC*. eLife 12, e79743, doi:10.7554/eLife.79743 (2023).

95 Menon, G. et al. Proximal termination generates a transcriptional state that determines the rate of establishment of Polycomb silencing. Mol. Cell 84, 2255–2271, doi:10.1016/j.molcel.2024.05.014 (2024).

96 Dean, C. & Howard, M. Graded versus ON/OFF control in quantitative gene expression and epigenetic memory. EMBO J. 45, 2869–2881, doi:10.1038/s44318-026-00750-y (2026).

97 Bergis-Ser, C. et al. LUMINIDEPENDENS orchestrates global transcriptional repression in *Arabidopsis*. Proc. Natl Acad. Sci. USA 122, e2527372122, doi:10.1073/pnas.2527372122 (2025).

98 Yang, M. et al. In vivo single-molecule analysis reveals *COOLAIR* RNA structural diversity. Nature 609, 394–399, doi:10.1038/s41586-022-05135-9 (2022).

99 Cao, X., Zhang, Y., Ding, Y. & Wan, Y. Identification of RNA structures and their roles in RNA functions. Nat. Rev. Mol. Cell Biol. 25, 784–801, doi:10.1038/s41580-024-00748-6 (2024).

100 Zhang, H. & Ding, Y. RNA structure: function and application in plant biology. Annu. Rev. Plant Biol. 76, 115–141, doi:10.1146/annurev-arplant-083123-055521 (2025).

101 Sun, Q., Csorba, T., Skourti-Stathaki, K., Proudfoot, N. J. & Dean, C. R-loop stabilization represses antisense transcription at the *Arabidopsis FLC* locus. Science 340, 619–621 (2013).

102 Clough, S. J. & Bent, A. F. Floral dip: a simplified method for *Agrobacterium*-mediated transformation of *Arabidopsis thaliana*. Plant J. 16, 735–743 (1998).

103 Livak, K. J. & Schmittgen, T. D. Analysis of relative gene expression data using real-time quantitative PCR and the 2^−ΔΔCT^ method. Methods 25, 402–408, doi:10.1006/meth.2001.1262 (2001).

104 Li, C. et al. Concerted genomic targeting of H3K27 demethylase REF6 and chromatin-remodeling ATPase BRM in *Arabidopsis*. Nature Genet. 48, 687–693, doi:10.1038/ng.3555 (2016).

105 Arora, D. et al. Establishment of proximity-dependent biotinylation approaches in different plant model systems. Plant Cell 32, 3388–3407, doi:10.1105/tpc.20.00235 (2020).

106 Liu, C. et al. A proxitome-RNA-capture approach reveals that processing bodies repress coregulated hub genes. Plant Cell 36, 559–584, doi:10.1093/plcell/koad288 (2024).

107 Jumper, J. et al. Highly accurate protein structure prediction with AlphaFold. Nature 596, 583–589, doi:10.1038/s41586-021-03819-2 (2021).

108 Abramson, J. et al. Accurate structure prediction of biomolecular interactions with AlphaFold 3. Nature 630, 493–500, doi:10.1038/s41586-024-07487-w (2024).

109 Zhao, L. et al. Integrative analysis of reference epigenomes in 20 rice varieties. Nat. Commun. 11, 2658, doi:10.1038/s41467-020-16457-5 (2020).

110 Zhang, Q. et al. aChIP is an efficient and sensitive ChIP-seq technique for economically important plant organs. Nat. Plants 10, 1317–1329, doi:10.1038/s41477-024-01743-7 (2024).

111 Yuan, L. et al. The transcriptional repressors VAL1 and VAL2 recruit PRC2 for genome-wide Polycomb silencing in *Arabidopsis*. Nucleic Acids Res. 49, 98–113, doi:10.1093/nar/gkaa1129 (2021).

112 Langmead, B. & Salzberg, S. L. Fast gapped-read alignment with Bowtie 2. Nat. Methods 9, 357–359, doi:10.1038/nmeth.1923 (2012).

113 Feng, J., Liu, T., Qin, B., Zhang, Y. & Liu, X. S. Identifying ChIP-seq enrichment using MACS. Nat. Protoc. 7, 1728–1740, doi:10.1038/nprot.2012.101 (2012).

114 Yu, G., Wang, L.-G. & He, Q.-Y. ChIPseeker: an R/Bioconductor package for ChIP peak annotation, comparison and visualization. Bioinformatics 31, 2382–2383, doi:10.1093/bioinformatics/btv145 (2015).

115 Ross-Innes, C. S. et al. Differential oestrogen receptor binding is associated with clinical outcome in breast cancer. Nature 481, 389–393, doi:10.1038/nature10730 (2012).

116 Thorvaldsdóttir, H., Robinson, J. T. & Mesirov, J. P. Integrative Genomics Viewer (IGV): high-performance genomics data visualization and exploration. Brief. Bioinform. 14, 178–192, doi:10.1093/bib/bbs017 (2013).

117 Ramírez, F. et al. deepTools2: a next generation web server for deep-sequencing data analysis. Nucleic Acids Res. 44, W160–W165, doi:10.1093/nar/gkw257 (2016).

118 Zhu, J., Liu, M., Liu, X. & Dong, Z. RNA polymerase II activity revealed by GRO-seq and pNET-seq in *Arabidopsis*. Nat. Plants 4, 1112–1123, doi:10.1038/s41477-018-0280-0 (2018).

119 Zhou, S. et al. Coupling of co-transcriptional splicing and 3′ end Pol II pausing during termination in *Arabidopsis*. Genome Biol. 24, 206, doi:10.1186/s13059-023-03050-4 (2023).

