## Supplemental Figures 1-10 for "Intragenic redistribution of RNA polymerase II licenses epigenetic silencing"

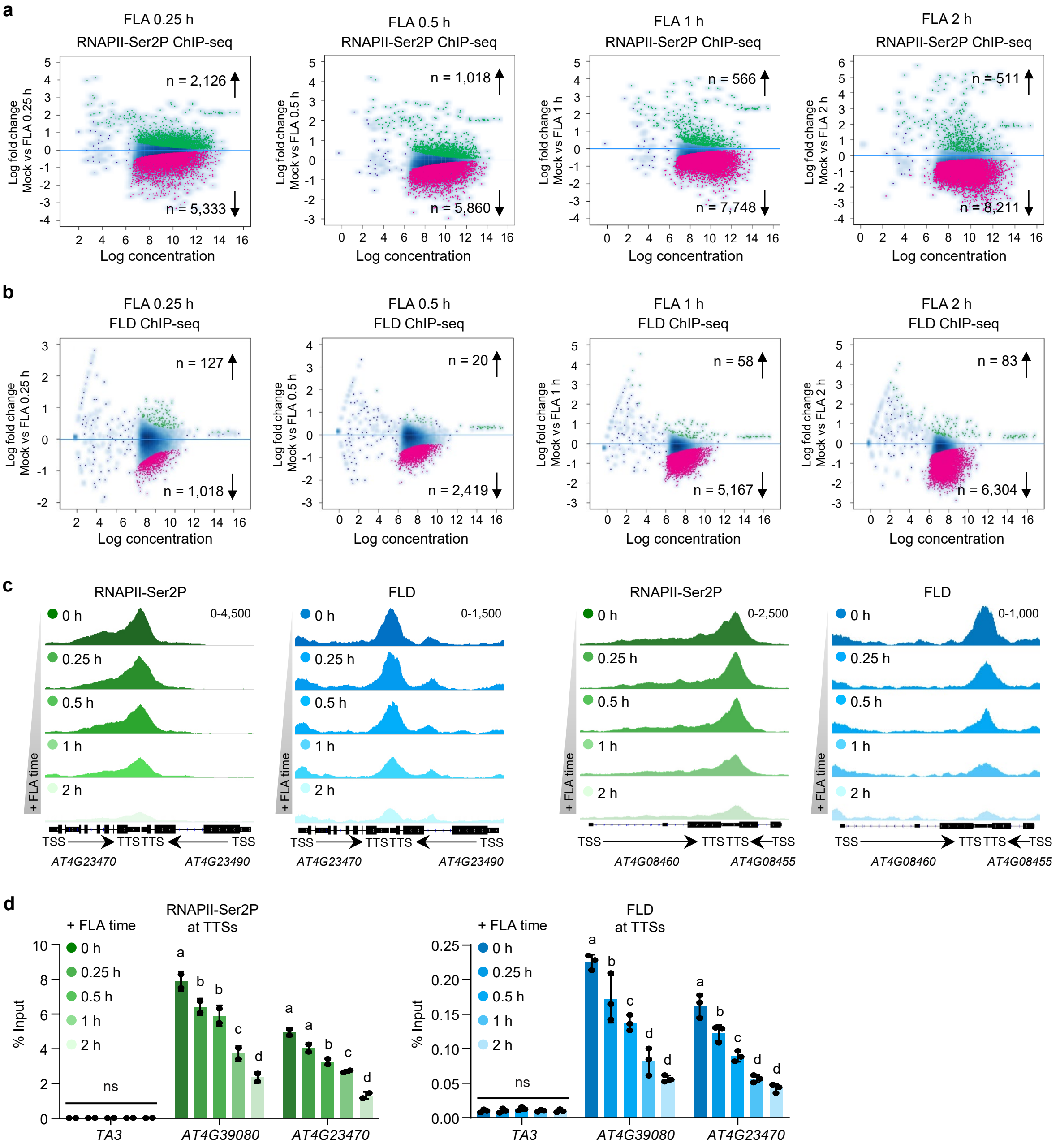


**Extended Data Fig. 1 | Temporal dynamics of RNAPII-Ser2P and FLD occupancy following flavopiridol treatment. a**, Scatter plots of RNAPII-Ser2P ChIP-seq occupancy comparing mock with flavopiridol (FLA) treatment for 0.25, 0.5, 1, and 2 h (left to right). Green, increased; magenta, decreased (FDR < 0.05). Total number of RNAPII-Ser2P peaks are 10,287. **b**, Scatter plots of FLD ChIP-seq occupancy comparing mock with flavopiridol (FLA) treatment for 0.25, 0.5, 1, and 2 h (left to right). Green, increased; magenta, decreased (FDR < 0.05). Total number of FLD peaks are 9,783. **c**, Integrative Genomics Viewer (IGV) tracks of RNAPII-Ser2P and FLD ChIP-seq signals at TTSs of representative RNAPII-Ser2P target genes after flavopiridol treatment. Black diagrams underneath indicate gene structures. **d**, ChIP-qPCR validation of RNAPII-Ser2P and FLD occupancy at the indicated loci across the flavopiridol time course. *TA3* serves as a negative control. Data are mean ± s.d. (*n* = 2 and 3 biological replicates for RNAPII-Ser2P and FLD, respectively). Different letters indicate significant differences (one-way ANOVA with Tukey's post hoc test).

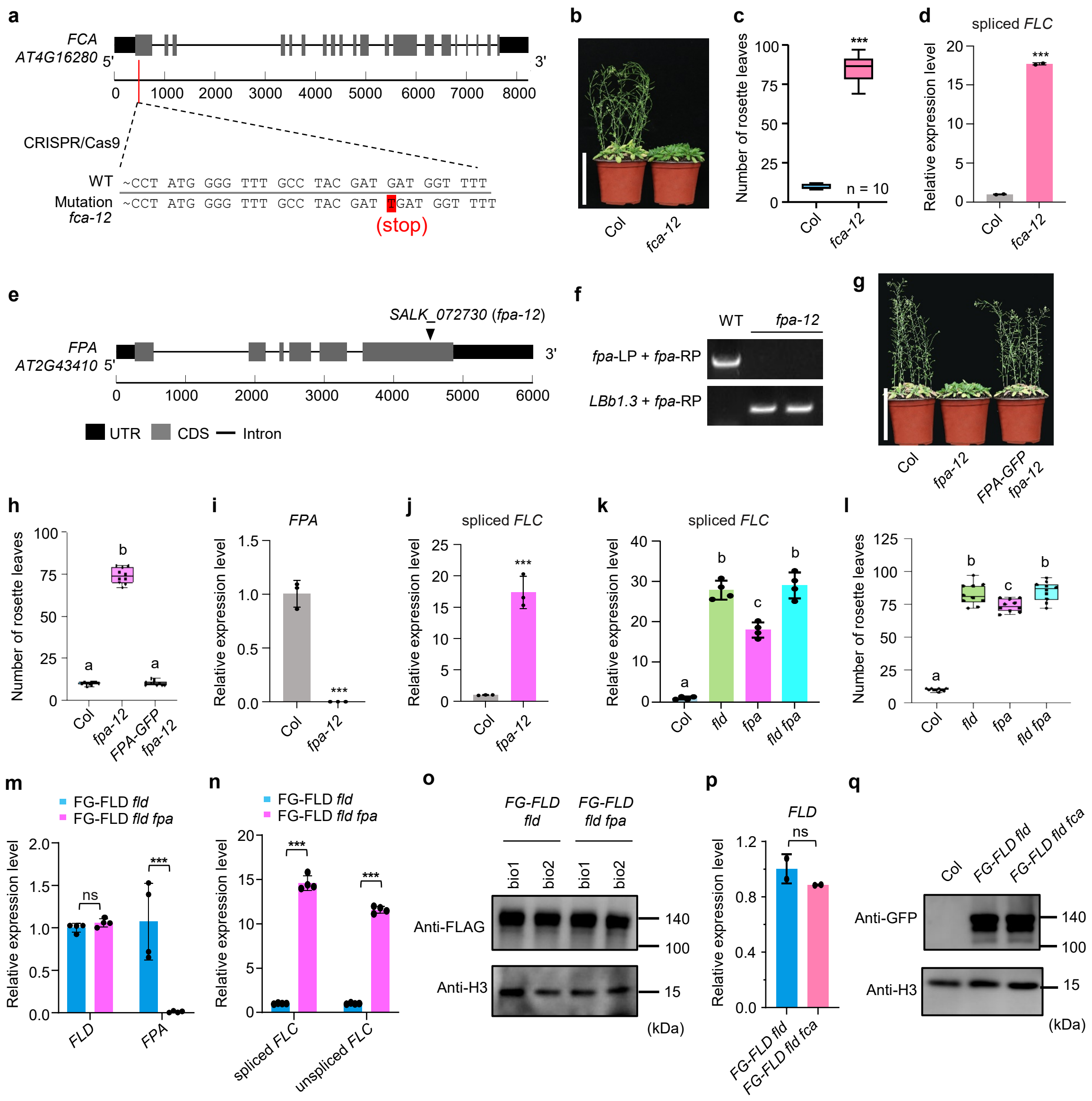
a
FCA
AT4G16280 5'
3'
0 1000 2000 3000 4000 5000 6000 7000 8000
CRISPR/Cas9
WT ~CCT ATG GGG TTT GCC TAC GAT GAT GGT TTT
Mutation fca-12 ~CCT ATG GGG TTT GCC TAC GAT TGAT GGT TTT
(stop)
b
Col
fca-12
c
Number of rosette leaves
n = 10
***
Col
fca-12
d
spliced FLC
Relative expression level
***
Col
fca-12
e
FPA
AT2G43410 5'
3'
SALK_072730 (fpa-12)
0 1000 2000 3000 4000 5000 6000
UTR CDS Intron
f
WT fpa-12
fpa-LP + fpa-RP
LBb1.3 + fpa-RP
g
Col
fpa-12
FPA-GFP fpa-12
h
Number of rosette leaves
a b a
Col
fpa-12
FPA-GFP fpa-12
i
FPA
Relative expression level
***
Col
fpa-12
j
spliced FLC
Relative expression level
***
Col
fpa-12
k
spliced FLC
Relative expression level
a b c b
Col
fld
fpa
fld fpa
l
Number of rosette leaves
a b c b
Col
fld
fpa
fld fpa
m
FG-FLD fld
FG-FLD fld fpa
Relative expression level
ns
***
FLD
FPA
n
FG-FLD fld
FG-FLD fld fpa
Relative expression level
***
***
spliced FLC
unspliced FLC
o
FG-FLD fld
FG-FLD fld fpa
bio1 bio2 bio1 bio2
Anti-FLAG
Anti-H3
140
100
15
(kDa)
p
FLD
ns
Relative expression level
FG-FLD fld
FG-FLD fld fca
q
Col
FG-FLD fld
FG-FLD fld fca
Anti-GFP
Anti-H3
140
100
15
(kDa)

**Extended Data Fig. 2 | Loss of FCA and FPA does not affect FLD protein abundance. a**, Schematic of *FCA* gene structure showing CRISPR/Cas9-induced mutation. Red box highlights the premature stop codon confirmed by Sanger sequencing. **b**, Flowering phenotype of Col and *fca-12* grown under long-day (LD) conditions for 5 weeks. Scale bar, 10 cm. **c**, Rosette leaf number at bolting. *n* = 10 plants. ****P* < 0.001, determined by two-tailed Student's *t*-test. **d**, Relative expression of spliced *FLC* in Col and *fca-12* at 7 DAG measured by RT-qPCR. Data are mean ± s.d. ****P* < 0.001, determined by two-tailed Student's *t*-test. **e**, Schematic of *FPA* gene structure showing T-DNA insertion site (black arrow) in *fpa-12*. **f**, Genotyping of *fpa-12* by three-primer PCR. WT serves as a control. **g**, Flowering phenotype of Col, *fpa-12* and *FPA-GFP fpa-12* grown under LD conditions for 5 weeks. Scale bar, 10 cm. **h**, Rosette leaf number at bolting. *n* = 10 plants. Different letters indicate significant differences (one-way ANOVA with Tukey's post hoc test). **i**, Relative expression of *FPA* in Col and *fpa-12* at 7 DAG. Data are mean ± s.d. (*n* = 3 biological replicates). ****P* < 0.001, determined by two-tailed Student's *t*-test. **j**, Relative expression of spliced *FLC* in Col and *fpa-12* at 7 DAG. Data are mean ± s.d. (*n* = 3 biological replicates). ****P* < 0.001, determined by two-tailed Student's *t*-test. **k**, Relative expression of spliced *FLC* in Col, *fld*, *fpa* and *fld fpa* at 7 DAG. Different letters indicate significant differences (one-way ANOVA with Tukey's post hoc test). Data are mean ± s.d. (*n* = 4 biological replicates). **l**, Rosette leaf number at bolting for indicated genotypes. *n* = 10 plants. Different letters indicate significant differences (one-way ANOVA with Tukey's post hoc test). **m**, Relative *FLD* transcript levels in *FG-FLD fld* and *FG-FLD fld fpa* backgrounds at 7 DAG. Data are mean ± s.d. (*n* = 4 biological replicates). ****P* < 0.001, determined by two-tailed Student's *t*-test. ns, not significant. **n**, Relative expression of spliced and unspliced *FLC* in *FG-FLD fld* and *FG-FLD fld fpa* backgrounds at 7 DAG. Data are mean ± s.d. (*n* = 4 biological replicates). ****P* < 0.001, determined by two-tailed Student's *t*-test. **o**, Immunoblot of FLD protein levels in *FG-FLD fld* and *FG-FLD fld fpa* backgrounds. H3 serves as a loading control. **p**, Relative *FLD* transcript levels in *FG-FLD fld* and *FG-FLD fld fca* backgrounds at 7 DAG. Data are mean ± s.d. (*n* = 2 biological replicates). ns, not significant, determined by two-tailed Student's *t*-test. **q**, Immunoblot showing FLD protein levels in *FG-FLD fld* and *FG-FLD fld fca* backgrounds. Histone H3 serves as a loading control.

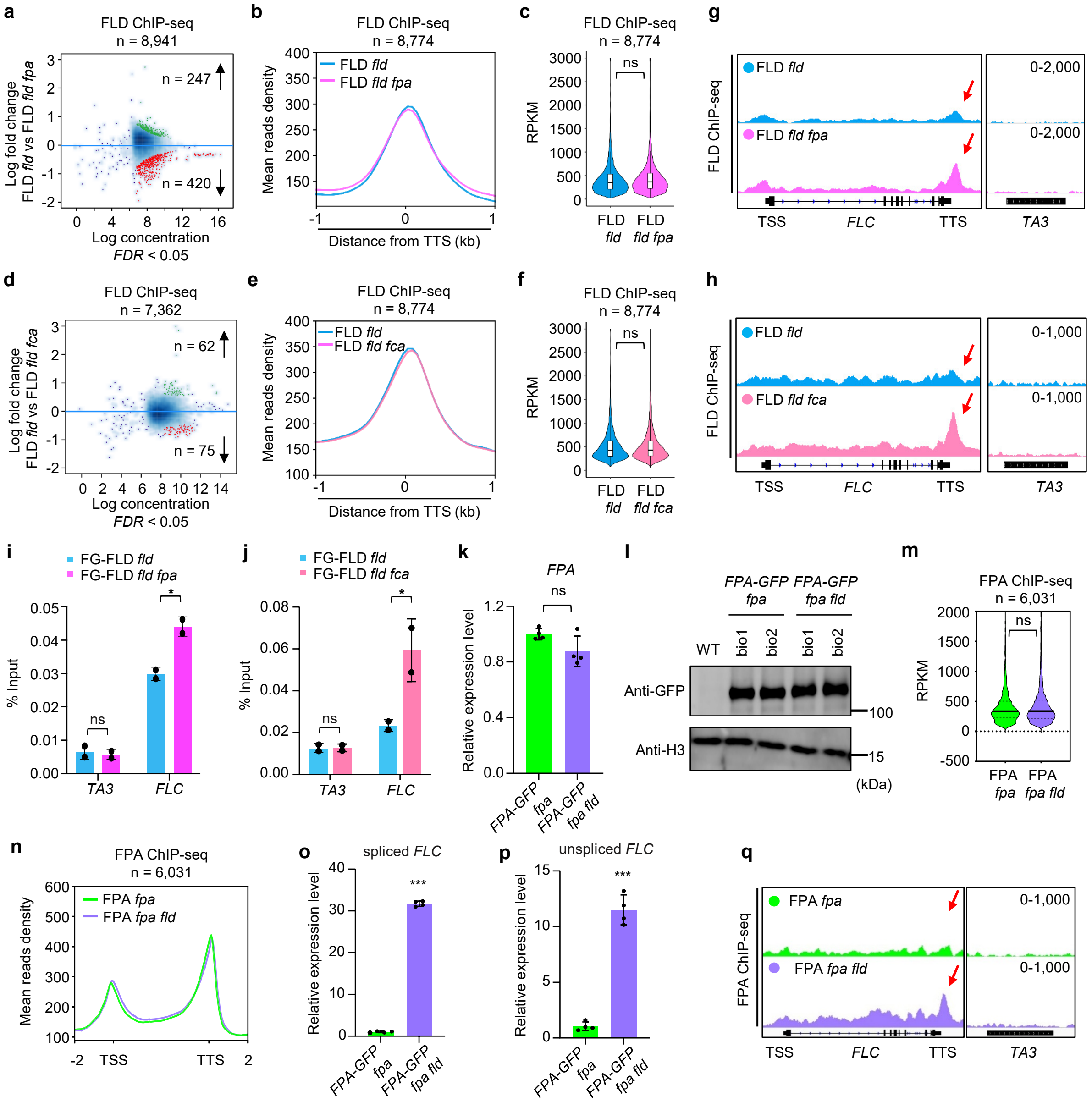
a
FLD ChIP-seq
n = 8,941
n = 247
n = 420
Log fold change
FLD fld vs FLD fld fpa
Log concentration
FDR < 0.05
b
FLD ChIP-seq
n = 8,774
FLD fld
FLD fld fpa
Mean reads density
Distance from TTS (kb)
c
FLD ChIP-seq
n = 8,774
ns
RPKM
FLD fld
FLD fld fpa
g
FLD ChIP-seq
FLD fld
FLD fld fpa
0-2,000
0-2,000
TSS
FLC
TTS
TA3
d
FLD ChIP-seq
n = 7,362
n = 62
n = 75
Log fold change
FLD fld vs FLD fld fca
Log concentration
FDR < 0.05
e
FLD ChIP-seq
n = 8,774
FLD fld
FLD fld fca
Mean reads density
Distance from TTS (kb)
f
FLD ChIP-seq
n = 8,774
ns
RPKM
FLD fld
FLD fld fca
h
FLD ChIP-seq
FLD fld
FLD fld fca
0-1,000
0-1,000
TSS
FLC
TTS
TA3
i
FG-FLD fld
FG-FLD fld fpa
% Input
ns
*
TA3
FLC
j
FG-FLD fld
FG-FLD fld fca
% Input
ns
*
TA3
FLC
k
FPA
ns
Relative expression level
FPA-GFP fpa
FPA-GFP fpa fld
l
FPA-GFP fpa
FPA-GFP fpa fld
WT
bio1
bio2
bio1
bio2
Anti-GFP
Anti-H3
100
15
(kDa)
m
FPA ChIP-seq
n = 6,031
ns
RPKM
FPA fpa
FPA fpa fld
n
FPA ChIP-seq
n = 6,031
FPA fpa
FPA fpa fld
Mean reads density
TSS
TTS
o
spliced FLC
***
Relative expression level
FPA-GFP fpa
FPA-GFP fpa fld
p
unspliced FLC
***
Relative expression level
FPA-GFP fpa
FPA-GFP fpa fld
q
FPA ChIP-seq
FPA fpa
FPA fpa fld
0-1,000
0-1,000
TSS
FLC
TTS
TA3

**Extended Data Fig. 3 | Transcriptional activity, not reciprocal recruitment, regulates FLD and FPA chromatin occupancy.** **a**, Scatter plot of FLD ChIP-seq occupancy in *fld* versus *fld fpa* backgrounds. Green, increased; red, decreased (FDR < 0.05). *n* = 8,941 peaks. **b**, Metagene plot of FLD ChIP-seq signal at the TTS (±1 kb) of FLD target genes in *fld* and *fld fpa* backgrounds. *n* = 8,774 genes. **c**, Violin plot of FLD ChIP-seq read density within ±500 bp of the TTS in *fld* and *fld fpa* backgrounds. *n* = 8,774 genes. ns, not significant; two-tailed Student's *t*-test. **d**, Scatter plot of FLD ChIP-seq occupancy in *fld* versus *fld fca* backgrounds. *n* = 7,362 peaks. **e**, Metagene plot of FLD ChIP-seq signal at the TTS (±1 kb) of FLD target genes in *fld* and *fld fca* backgrounds. *n* = 8,774 genes. **f**, Violin plot of FLD ChIP-seq read density within ±500 bp of the TTS in *fld* and *fld fca* backgrounds. *n* = 8,774 genes. ns, not significant; two-tailed Student's *t*-test. **g**, FLD ChIP-seq tracks at *FLC* in *fld* and *fld fpa* backgrounds. Red arrows indicate FLD peaks. *TA3* serves as a negative control. **h**, FLD ChIP-seq tracks at *FLC* in *fld* and *fld fca* backgrounds. Red arrows indicate FLD peaks. *TA3* serves as a negative control. **i**,**j**, ChIP-qPCR validation of FLD occupancy at *FLC* and *TA3* (negative control) in *fld* versus *fld fpa* (**i**) and *fld* versus *fld fca* (**j**) backgrounds. Data are mean ± s.d. (*n* = 2 biological replicates). **P* < 0.05, determined by two-tailed Student's *t*-test; ns, not significant. **k**, Relative *FPA* transcript levels in FPA-GFP *fpa* and FPA-GFP *fpa fld* backgrounds at 7 DAG. Data are mean ± s.d. (*n* = 4 biological replicates). ns, not significant; two-tailed Student's *t*-test. **l**, Immunoblot showing FPA-GFP protein levels in *fpa* and *fpa fld* backgrounds. H3 serves as a loading control. **m**, Violin plot of FPA ChIP-seq read density within ±1 kb of the TTS in *FPA-GFP fpa* and *FPA-GFP fpa fld* backgrounds. *n* = 6,031 genes. ns, not significant; two-tailed Student's *t*-test. **n**, Metagene plot of FPA ChIP-seq signal at FPA target genes in *fpa* and *fpa fld* backgrounds. *n* = 6,031 genes. **o**,**p**, Relative expression of spliced *FLC* (**o**) and unspliced *FLC* (**p**) in *FPA-GFP fpa* and *FPA-GFP fpa fld* backgrounds at 7 DAG. Data are mean ± s.d. (*n* = 4 biological replicates). ****P* < 0.001, determined by two-tailed Student's *t*-test. **q**, IGV tracks of FPA ChIP-seq signals at *FLC* in *FPA-GFP fpa* and *FPA-GFP fpa fld* backgrounds. *TA3* serves as a negative control. The black diagrams underneath indicate gene structures.

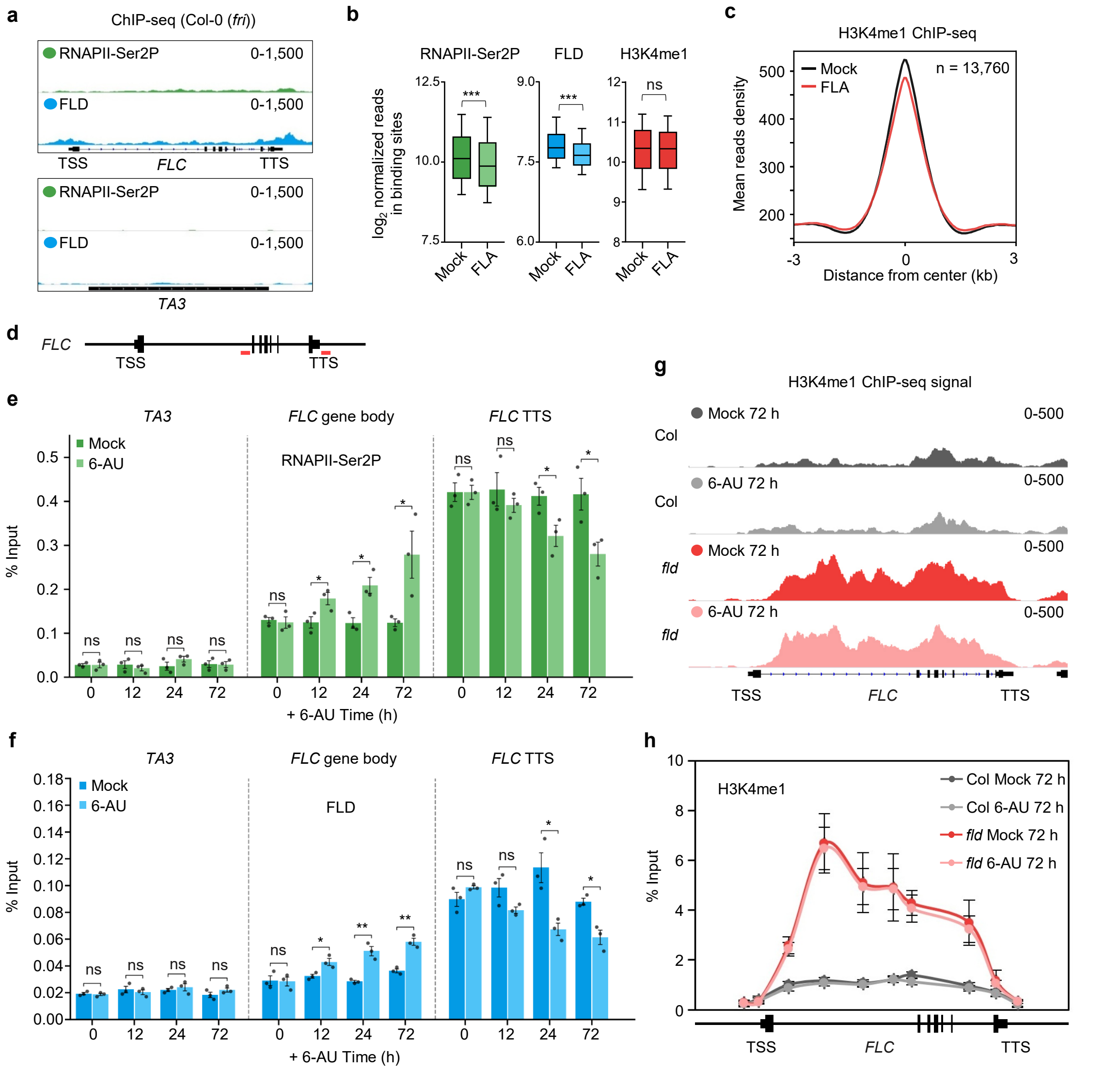


**Extended Data Fig. 4 | Supporting data for flavopiridol and 6-AU treatments at the *FLC* locus.** **a**, ChIP-seq tracks of RNAPII-Ser2P and FLD at the *FLC* and *TA3* loci in Col. **b**, Box plots of $log_2$-normalized RNAPII-Ser2P, FLD, and H3K4me1 ChIP-seq reads at binding sites after mock or flavopiridol (FLA) treatment. Statistical significance was determined by two-tailed Student's *t*-test; ****P* < 0.001; ns, not significant. **c**, Metagene plot of H3K4me1 ChIP-seq signal at genome-wide peaks (*n* = 13,760) after mock or FLA treatment. **d**, Schematic of the *FLC* gene structure. Red lines indicate the positions of the ChIP-qPCR primer sets used in **e** and **f**: *FLC* gene body and *FLC* TTS. **e**,**f**, ChIP-qPCR validation of RNAPII-Ser2P (**e**) and FLD (**f**) occupancy at *TA3* (negative control), *FLC* gene body and *FLC* TTS following mock or 6-AU treatment for 0, 12, 24, and 72 h. Data are mean ± s.d. (*n* = 3 biological replicates). **P* < 0.05; ***P* < 0.01; ns, not significant. **g**, H3K4me1 ChIP-seq tracks at the *FLC* locus after 72 h of mock or 6-AU treatment in Col and *fld* backgrounds. **h**, ChIP-qPCR validation of H3K4me1 levels across *FLC* after 72 h of mock or 6-AU treatment in Col and *fld* backgrounds. Data are mean ± s.d. (n = 3 biological replicates).

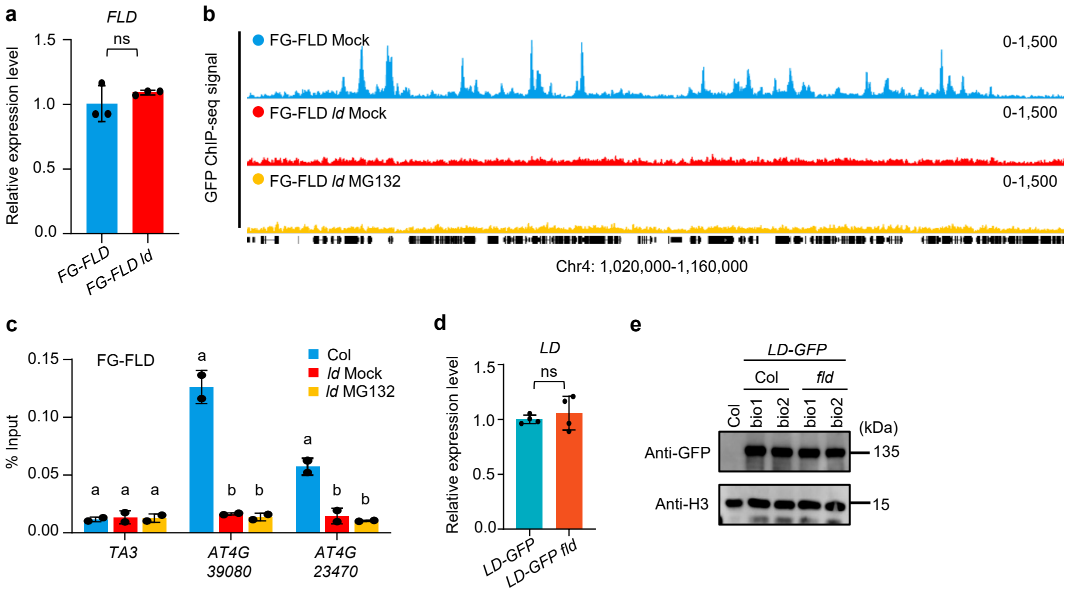


**Extended Data Fig. 5 | FLD chromatin occupancy depends on LD. a**, Relative *FLD* transcript levels in Col and *ld* backgrounds measured by RT-qPCR at 7 DAG. Data are mean ± s.d. ($n$ = 3 biological replicates). ns, not significant; two-tailed Student's $t$-test. **b**, FLD ChIP-seq tracks across a representative genomic region (Chr4: 1,020,000–1,160,000). **c**, ChIP-qPCR validation of FLD occupancy at the indicated loci in Col, *ld*, and MG132-treated *ld* seedlings, performed as independent validation of the corresponding ChIP-seq results. *TA3* serves as a negative control. Data are mean ± s.d. ($n$ = 2 biological replicates). Different letters indicate significant differences (one-way ANOVA with Tukey's post hoc test). **d**, Relative *LD* transcript levels in Col and *fld* backgrounds measured by RT-qPCR at 7 DAG. Data are mean ± s.d. ($n$ = 4 biological replicates). ns, not significant; two-tailed Student's $t$-test. **e**, Immunoblot showing LD-GFP protein levels in Col and *fld* backgrounds. H3 serves as a loading control.

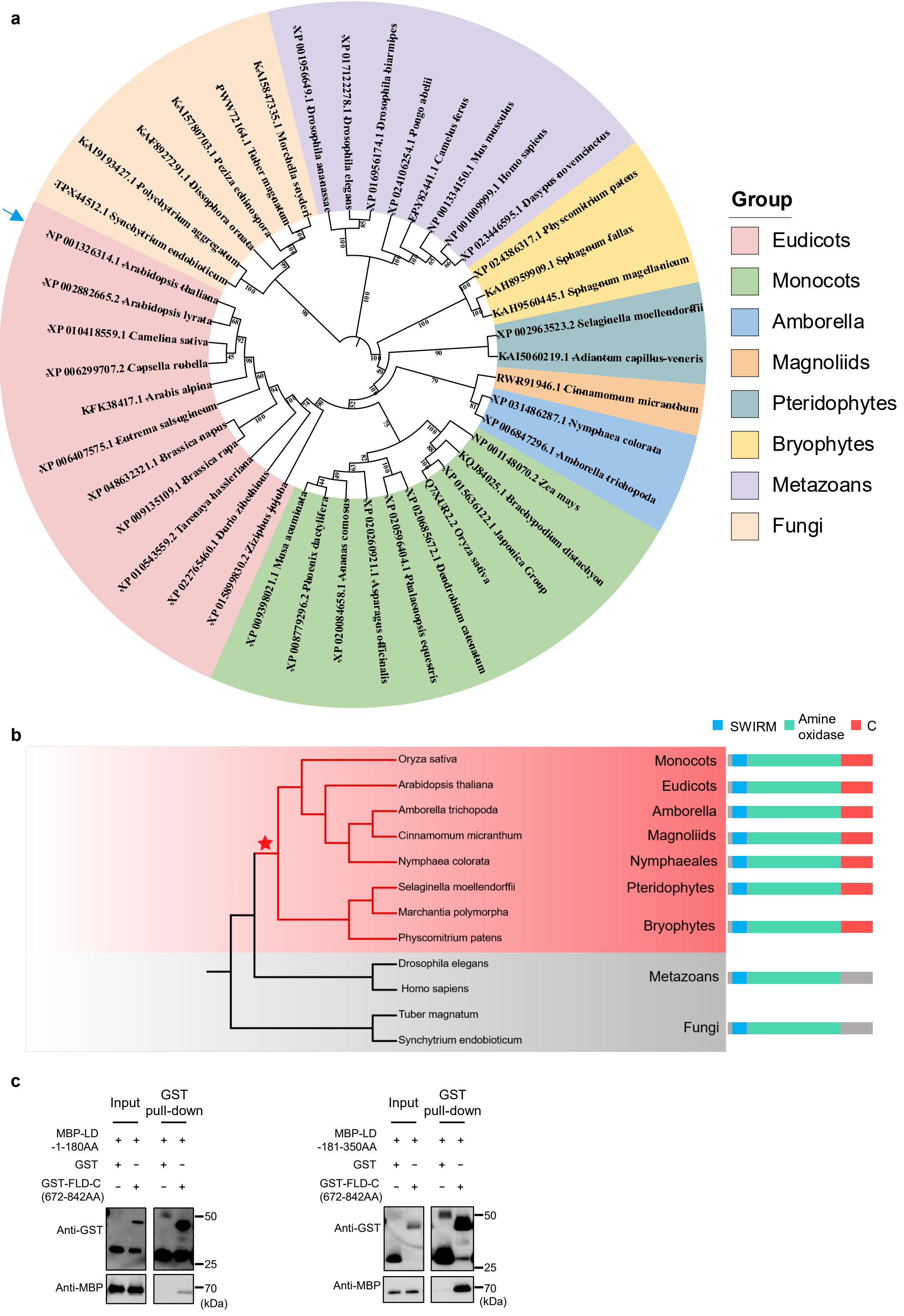
a
Group
Eudicots
Monocots
Amborella
Magnoliids
Pteridophytes
Bryophytes
Metazoans
Fungi
XP 001956649.1 Drosophila ananassae
XP 017122278.1 Drosophila elegans
XP 016956174.1 Drosophila biarmipes
XP 024106254.1 Pongo abelii
EPY82441.1 Camelus ferus
NP 001334150.1 Mus musculus
NP 001009999.1 Homo sapiens
XP 023446595.1 Dasypus novemcinctus
XP 024386317.1 Physcomitrium patens
KAH8959909.1 Sphagnum fallax
KAH9560445.1 Sphagnum magellanicum
XP 002963523.2 Selaginella moellendorffii
KAI5060219.1 Adiantum capillus-veneris
RWR91946.1 Cinnamomum micranthum
XP 031486287.1 Nymphaea colorata
XP 006847296.1 Amborella trichopoda
NP 001148070.2 Zea mays
KQJ84025.1 Brachypodium distachyon
XP 015636122.1 Japonica Group
Q7XUR2.2 Oryza sativa
XP 020685672.1 Dendrobium catenatum
XP 020590404.1 Phalaenopsis equestris
XP 020260921.1 Asparagus officinalis
XP 020084658.1 Ananas comosus
XP 008779296.2 Phoenix dactylifera
XP 009398021.1 Musa acuminata
XP 015899830.2 Ziziphus jujuba
XP 022765460.1 Durio zibethinus
XP 010543559.2 Tarenaya hassleriana
XP 009135109.1 Brassica rapa
XP 048632321.1 Brassica napus
XP 006407575.1 Eutrema salsugineum
KFK38417.1 Arabis alpina
XP 006299707.2 Capsella rubella
XP 010418559.1 Camelina sativa
XP 002882665.2 Arabidopsis lyrata
NP 001326314.1 Arabidopsis thaliana
TPX44512.1 Synchytrium endobioticum
KAI9193427.1 Polychytrium aggregatum
KAF8927291.1 Dissophora ornata
KAI5780703.1 Peziza echinospora
PWW72164.1 Tuber magnatum
KAI5847335.1 Morchella snyderi
b
SWIRM
Amine oxidase
C
Oryza sativa
Arabidopsis thaliana
Amborella trichopoda
Cinnamomum micranthum
Nymphaea colorata
Selaginella moellendorffii
Marchantia polymorpha
Physcomitrium patens
Drosophila elegans
Homo sapiens
Tuber magnatum
Synchytrium endobioticum
Monocots
Eudicots
Amborella
Magnoliids
Nymphaeales
Pteridophytes
Bryophytes
Metazoans
Fungi
c
Input
GST pull-down
MBP-LD -1-180AA
MBP-LD -181-350AA
GST
GST-FLD-C (672-842AA)
Anti-GST
Anti-MBP
50
25
70
(kDa)

**Extended Data Fig. 6 | Phylogenetic analysis of the FLD C-terminal region and mapping of its interaction with LD. a**, Phylogenetic tree of FLD and homologous LSD1-family proteins across eukaryotes. The tree was constructed using full-length amino acid sequences. Blue arrow points the Arabidopsis thaliana FLD. **b**, Domain architecture of FLD homologues across representative species. The SWIRM (blue), amine oxidase (green) and the C-terminal region (red) are indicated. The red star marks the inferred emergence of the C-terminal region in land plants. Taxonomic groups include monocots, eudicots, *Amborella*, magnoliids, Nymphaeales, pteridophytes, bryophytes, metazoans, and fungi. **c**, GST pull-down assays mapping the FLD-C–LD interaction. Left, GST-FLD-C (amino acids 672–842) or GST alone with MBP-LD (amino acids 1–180, containing HOX domain). Right, GST-FLD-C or GST alone with MBP-LD (amino acids 181–350, containing TFIIS domain). Anti-GST and anti-MBP immunoblots of the input and pull-down fractions are shown.

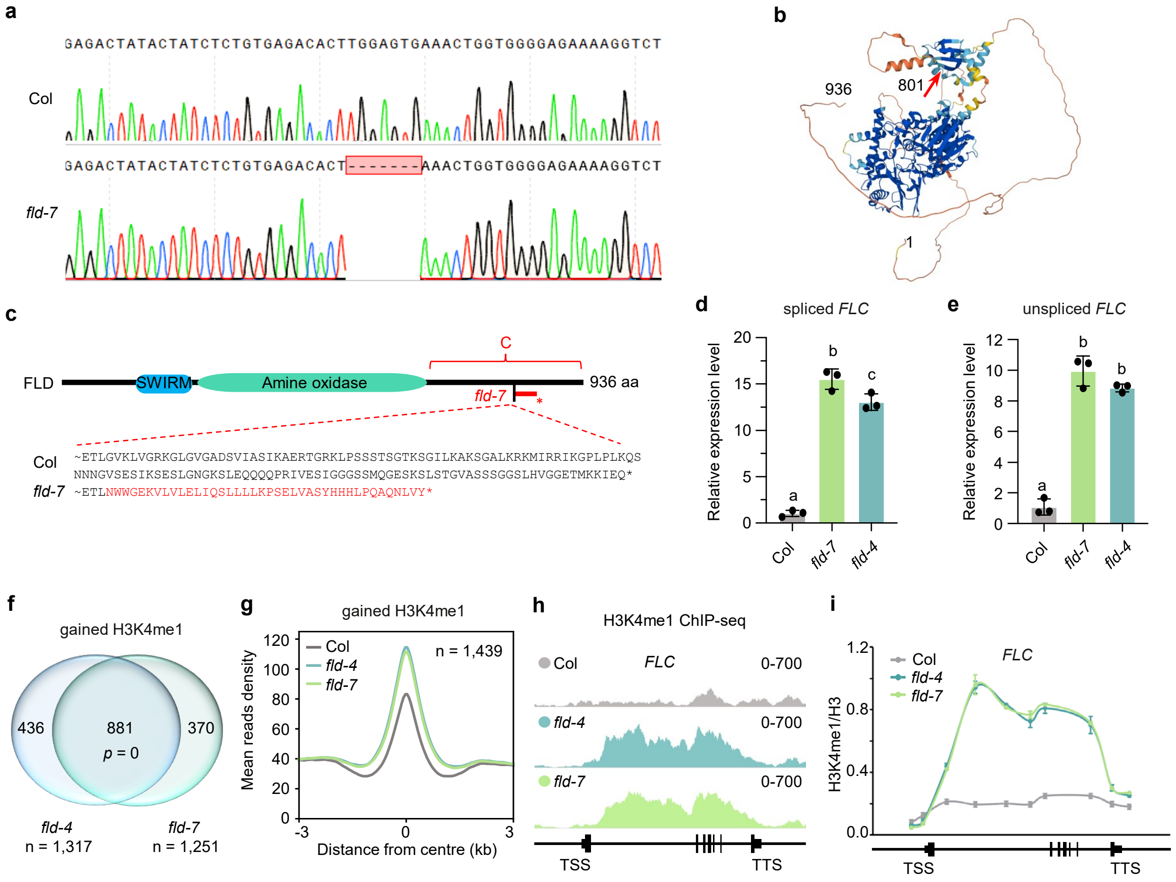


**Extended Data Fig. 7 | Molecular and functional characterization of the *fld-7* allele. a**, Sanger sequencing chromatograms; the red box highlights the 7-bp deletion. **b**, AlphaFold-predicted FLD protein structure; the red arrow indicates the frameshift position in *fld-7* (residue 801). **c**, Schematic of FLD domain structure showing the position of the *fld-7* mutation. The WT sequence and frameshift caused by the 7-bp deletion are shown below. **d**,**e**, Relative expression of spliced *FLC* (**d**) and unspliced *FLC* (**e**) in Col and *fld* mutants at 7 DAG, measured by RT-qPCR. Data are mean ± s.d. (*n* = 3 biological replicates). Different letters indicate significant differences (one-way ANOVA with Tukey's post hoc test). **f**, Venn diagram showing overlap of H3K4me1-gained genes between *fld-4* (*n* = 1,317) and *fld-7* (*n* = 1,251). *P* value was calculated using a hypergeometric test. **g**, Metagene plot of H3K4me1 ChIP-seq signals at gained peaks (*n* = 1,439) in Col, *fld-4*, and *fld-7*. **h**, H3K4me1 ChIP-seq tracks at *FLC* in Col, *fld-4*, and *fld-7*. **i**, ChIP-qPCR validation of H3K4me1 levels at the *FLC* locus in Col, *fld-4*, and *fld-7*. Data are mean ± s.d. (*n* = 3 biological replicates).

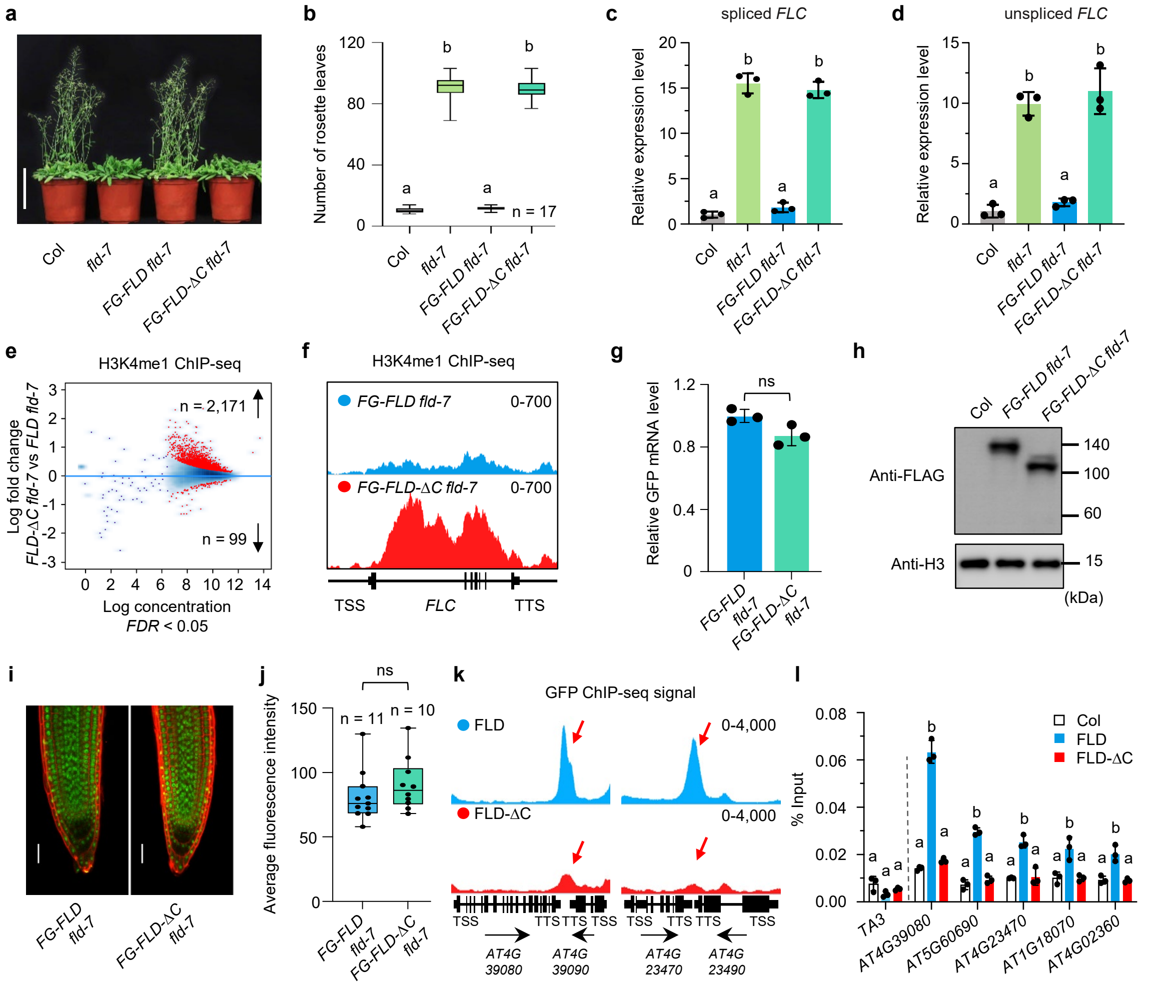


**Extended Data Fig. 8 | The C-terminal region of FLD is required for FLD chromatin engagement and H3K4me1 demethylation.** **a**, Flowering phenotypes of indicated genotypes. Scale bar, 10 cm. **b**, Rosette leaf number at bolting. *n* = 17 plants. Different letters indicate significant differences (one-way ANOVA with Tukey's post hoc test). **c**,**d**, Relative expression of spliced *FLC* (**c**) and unspliced *FLC* (**d**) at 7 DAG. Data are mean ± s.d. (*n* = 3 biological replicates). Different letters indicate significant differences (one-way ANOVA with Tukey's post hoc test). **e**, Scatter plot of H3K4me1 ChIP-seq signal in *FG-FLD-ΔC fld-7* versus *FG-FLD fld-7* plants. Red, differentially occupied peaks (FDR < 0.05). **f**, H3K4me1 ChIP-seq tracks at *FLC* in *FG-FLD fld* and *FG-FLD-ΔC fld* backgrounds. **g**, Relative *GFP-FLD* transcript levels in *FG-FLD fld-7* and *FG-FLD-ΔC fld-7* at 7 DAG, measured by RT-qPCR. Data are mean ± s.d. (*n* = 3 biological replicates). Different letters indicate significant differences. **h**, Immunoblot showing FG-FLD and FG-FLD-ΔC protein levels. H3 serves as a loading control. **i**, Confocal images of root tips showing nuclear localization of FG-FLD and FG-FLD-ΔC proteins. Red, propidium iodide staining. Scale bar, 50 μm. **j**, Quantification of GFP fluorescence intensity. *n*, number of roots analysed. ns, not significant. **k**, FG-FLD and FG-FLD-ΔC ChIP-seq tracks at TTSs of representative convergently transcribed genes. **l**, ChIP-qPCR validation at the indicated loci. *TA3* serves as a negative control. Data are mean ± s.d. (*n* = 3 biological replicates). Different letters indicate significant differences (one-way ANOVA with Tukey's post hoc test).

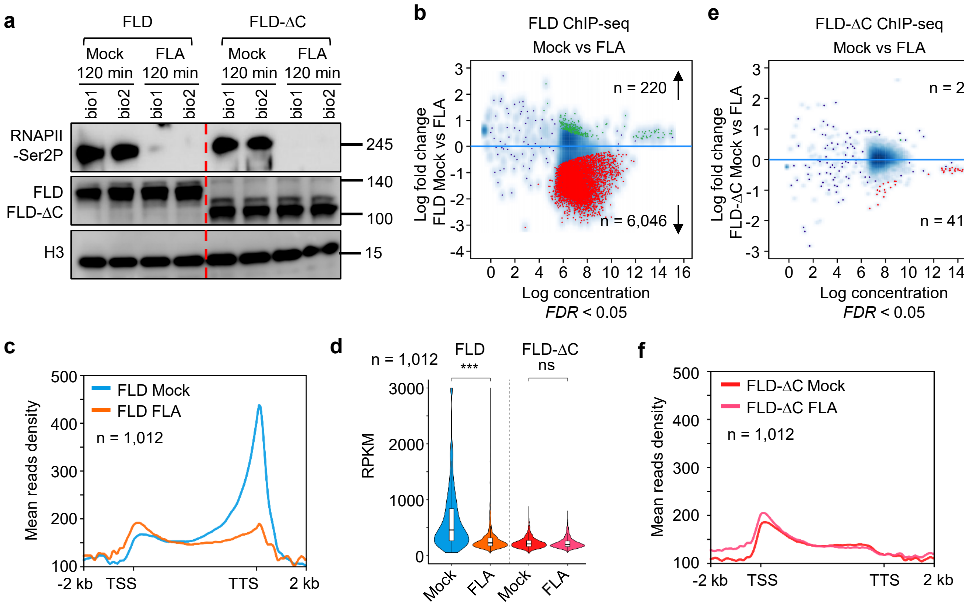


**Extended Data Fig. 9 | The C-terminal region of FLD mediates RNAPII-dependent chromatin engagement. a**, Immunoblot showing RNAPII-Ser2P, FLD, and FLD-ΔC protein levels after 2 h of mock or flavopiridol (FLA) treatment in 12-day-old seedlings. H3 serves as a loading control. **b**, Scatter plots of FLD chromatin occupancy comparing mock and FLA treatments. Green, increased; red, decreased (FDR < 0.05). **c**, Metagene plots of FLD ChIP-seq signal after mock or FLA treatment at RNAPII-Ser2P target genes forming convergent gene pairs ($n$ = 1,012). **d**, Violin plot of ChIP-seq read density for FLD and FLD-ΔC under mock and FLA conditions. ***$P$ < 0.001; ns, not significant; two-tailed Student's $t$-test. **e**, Scatter plots of FLD-ΔC chromatin occupancy comparing mock and FLA treatments. Green, increased; red, decreased (FDR < 0.05). **f**, Metagene plots of FLD-ΔC ChIP-seq signal after mock or FLA treatment at RNAPII-Ser2P target genes forming convergent gene pairs ($n$ = 1,012).

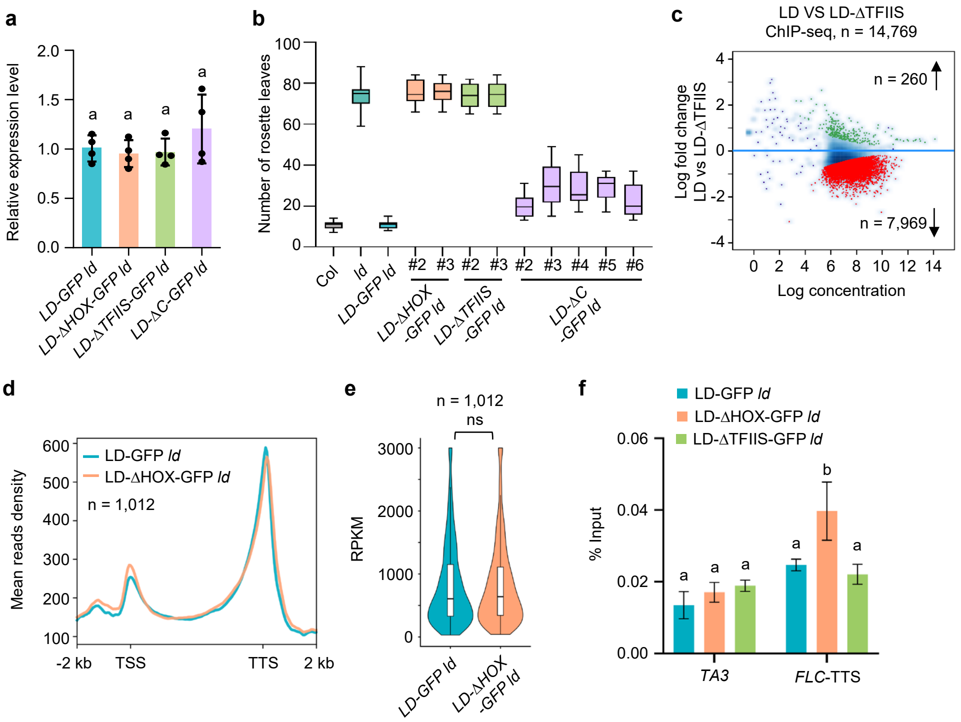


**Extended Data Fig. 10 | Characterization of LD domain-deletion variants. a**, Relative expression of *LD-GFP*, *LD-ΔHOX-GFP*, *LD-ΔTFIIS-GFP*, and *LD-ΔC-GFP* transgenes in the *ld* background, measured by RT-qPCR. Data are mean ± s.d. ($n$ = 4 biological replicates). Different letters indicate significant differences. **b**, Rosette leaf number at bolting for independent transgenic lines of *LD-GFP ld*, *LD-ΔHOX-GFP ld*, *LD-ΔTFIIS-GFP ld*, and *LD-ΔC-GFP ld*. Line numbers are indicated. **c**, Scatter plot comparing LD-GFP and LD-ΔTFIIS-GFP ChIP-seq occupancy in the *ld* background. Differentially occupied peaks (FDR < 0.05) are highlighted: decreased (red, $n$ = 7,969) and increased (green, $n$ = 260). $n$ = 14,769 total peaks. **d**,**e**, Metagene plot (**d**) and violin plot (**e**) of LD-GFP and LD-ΔHOX-GFP ChIP-seq signals at RNAPII-Ser2P target genes forming convergent gene pairs (as in Fig. 1D) ($n$ = 1,012). ns, not significant; two-tailed Student's $t$-test. **f**, ChIP-qPCR showing LD, LD-ΔHOX, and LD-ΔTFIIS occupancy at the *FLC* TTS in *ld* mutant background. *TA3* serves as a negative control locus. Data are mean ± s.d. Different letters indicate significant differences (one-way ANOVA with Tukey's post hoc test).
